# Dual contributions of Xrp1 to genome integrity through the DNA damage response and cell competition

**DOI:** 10.1101/2022.06.06.494998

**Authors:** Chaitali Khan, Nasser M Rusan, Nicholas E Baker

## Abstract

Model organisms may help understand how p53 suppresses tumorigenesis in mammals. In *Drosophila*, the primary transcriptional target of p53 is the gene encoding the bZip AT-hook protein Xrp1, which is another transcription factor. We report that Xrp1 mediated multiple functions of p53 in the DNA damage response (DDR), contributing to p53-dependent gene transcription and DNA damage-induced apoptosis. In addition to this role as a p53 effector, a p53-independent role for Xrp1 in cell competition has been described. Cell competition can remove cells whose genome has been altered by DNA damage and repair. During cell competition, Xrp1 is induced by RpS12, which acts as a sensor of defective ribosome biogenesis. RpS12-dependent cell competition began as the DDR wound down, and was even more prominent if p53 function was reduced. Such p53 inhibition resulted in persistence of DNA damage revealed by γH2Av accumulation. Thus, Xrp1 limited the accumulation of abnormal cells resulting from genotoxicity through both the acute, p53-dependent DDR, and also from subsequent cell competition that removes cells where DNA repair did not restore the normal genome. Both these processes might contribute to the tumor suppressor function of p53 in mammals.

## INTRODUCTION

Maintaining genome integrity is essential not only to preserve genetic information in the germline but also for somatic cells. Defects in DNA repair are associated with elevated cancer incidences, and defects in mitotic fidelity occur in aging (Tiwari & Wilson, 2019; Vijg, 2021; Hopkins et al., 2022; Nemeth & Szuts, 2024). Most cancer cells are aneuploid, and both mouse models and human genetic diseases that predispose to aneuploidy are cancer prone (Hanks et al., 2004; Baker et al., 2009; Baker & van Deursen, 2010). The most important human tumor suppressor, p53, is well known as a master regulator of the DNA damage response (DDR) that also maintains the integrity of somatic tissues by the apoptotic elimination of cells with damaged DNA (Levine et al., 2006; Zhang et al., 2011). P53 also has other functions, and it has been questioned whether the DNA damage response is, in fact, responsible for its tumor suppressor function (Kaiser & Attardi, 2018; Boutelle & Attardi, 2021). In addition to apoptosis as a cell-autonomous response to DNA damage, a process of cell competition has also been described in which damaged cells can be removed only when they are less fit than their neighbors (Baker, 2020; Baker & Montagna, 2022; Kiparaki & Baker, 2023; Khandekar & Ellis, 2024; Nagata & Igaki, 2024). Cell competition is able to remove cells with damage that would be tolerated by cell-autonomous mechanisms. In *Drosophila*, cell competition occurs independently of p53 (Kale et al., 2015), but in mammals, p53 activities too low to cause apoptosis by cell-autonomous mechanisms can trigger competitive cell removal when compared with less affected surrounding cells (Bondar & Medzhitov, 2010a; Wagstaff et al., 2016; Zhang et al., 2017; Fernandez-Antoran et al., 2019).

In *Drosophila*, the *Xrp1* gene is a common link between the DNA damage response and cell competition. Xrp1 was first described as a transcriptional target of p53, encodes a *Drosophila* bZip domain protein, and is transcriptionally induced by p53 when *Drosophila* embryos are irradiated (Brodsky et al., 2004; Akdemir et al., 2007). Flies lacking wild-type Xrp1 exhibit increased sensitivity to irradiation. Specifically, they show increased loss of heterozygosity in an assay that quantifies patches of cells expressing the recessive *mwh* phenotype in otherwise *mwh^+/-^* individuals, indicative of somatic loss of the *mwh^+^* allele (Akdemir et al., 2007). This loss of heterozygosity following irradiation is consistent with a role for Xrp1 in the DDR, regulated by p53.

Xrp1 was later identified as a central regulator of the cell competition of ‘Minute’ (Ribosome protein gene heterozygous mutant) cells in imaginal discs (Lee et al., 2016; Baillon et al., 2018; Lee et al., 2018b; Brown et al., 2021). Cell competition was first described as a phenomenon affecting cells heterozygous for mutations in genes encoding Ribosomal proteins (Rp) (Morata & Ripoll, 1975). Most Rp genes are essential, so *Rp* mutations are generally homozygously lethal, whereas *Rp^+/-^* heterozygotes survive with a developmental delay, thin bristles, and some other traits (Bridges & Morgan, 1923; Marygold et al., 2007). In contrast to whole *Rp^+/-^* animals, individual *Rp^+/-^* cells or clones are eliminated from many tissues when wild-type cells are also present, suffering an enhanced rate of apoptosis (Morata & Ripoll, 1975; Simpson, 1979; Moreno et al., 2002; Li & Baker, 2007). The Xrp1 protein is expressed at only low levels in wild-type imaginal discs, but Xrp1 protein is robustly induced in *Rp^+/-^* cells, where it is required for transcriptional changes in *Rp^+/-^* wing discs and for many aspects of the *Rp^+/-^* phenotype, including both developmental delay and cell competition (Baillon et al., 2018; Lee et al., 2018b; Boulan et al., 2019). Xrp1 binds to DNA as a heterodimer with another protein, Irbp18. Unlike Xrp1, Irbp18 is abundantly expressed in wild-type imaginal disc cells (Reinke et al., 2013; Francis et al., 2016; Blanco et al., 2020). The Xrp1/Irbp18 heterodimer activates transcription by binding to a sequence motif identified from both in vitro studies and ChIP-Seq *in vivo* (Zhu et al., 2011; Baillon et al., 2018; Kiparaki et al., 2022). Like Xrp1, Irbp18 is also required for cell competition, suggesting that transcriptional targets of the Xrp1/Irbp18 heterodimer are involved (Blanco et al., 2020).

Unlike Xrp1 expression following irradiation, Xrp1 induction in *Rp^+/-^* cells does not require p53, and p53 is also not needed for the competitive elimination of *Rp^+/-^* cells from mosaics (Kale et al., 2015; Lee et al., 2018b). In *Rp^+/-^* cells, Xrp1 expression is induced by a particular Rp, RpS12, which is necessary for cell competition (Kale et al., 2018; Lee et al., 2018b; Ji et al., 2019). Mutations in other *Rp* genes, and other ribosome biogenesis defects such as diminished rRNA synthesis, behave as though they increase RpS12 function (Baker & Kale, 2016; Ji et al., 2019; Kiparaki et al., 2022). RpS12 activates Xrp1 expression through alternative splicing of the Xrp1 mRNA (Kakemura et al., 2025; Potiri et al., 2025). Other cellular stresses can also activate Xrp1 expression, at least some independently of both p53 and RpS12 (Baumgartner et al., 2021; Brown et al., 2021; Ochi et al., 2021; Recasens-Alvarez et al., 2021; Kiparaki et al., 2022; Kumar & Baker, 2022; Kiparaki & Baker, 2023; Shekhar et al., 2025).

In *Drosophila*, cell competition is part of the response to aneuploidy(Titen & Golic, 2008; McNamee & Brodsky, 2009; Ji et al., 2021a; Baker & Montagna, 2022). Because the genes encoding 80 Rp are distributed around the chromosomes, loss of chromosome segments constituting more than 1.2% of the genome (on average) typically leads to haploinsufficiency for one *Rp* gene or another by removing one copy of that locus. This leads to the elimination of cells bearing many segmental monosomies by the same cell competition pathway that affects *Rp^+/-^* point mutant cells (Ji et al., 2021b). Accordingly, *Drosophila* lacking cell competition can accumulate cells with certain kinds of genetic damage, not because they experience any defect in DNA repair, but because some abnormal genotypes that would ordinarily be eliminated by cell competition survive when cell competition is inhibited. Adult flies with a mutation in RpS12 that prevents cell competition exhibit an increased frequency of sporadic cells exhibiting the typical *Rp^+/-^*mutant thin bristle phenotype, whether occurring spontaneously or after irradiation. Many of these cells likely harbor chromosome deletions that affect multiple genes in addition to one of the *Rp* loci (Ji et al., 2021b).

Because Xrp1 has been implicated in two processes that impact genome integrity, we sought here to define the respective roles more precisely. Loss of heterozygosity after irradiation of Xrp1 mutants could either arise through defects in p53-dependent DNA repair or apoptosis, or due to failure to outcompete cells bearing irradiation-induced segmental aneuploidies. We now report that Xrp1 is indeed important for the DDR as well as for cell competition, mediating the expression of most of the p53-target genes and contributing to p53-induced apoptosis and the repair or removal of DNA damage. If cells with unrepaired DNA and damaged genomes accumulate after DNA repair, many of these are subsequently removed by p53-independent, Xrp1-dependent cell competition. Thus, Xrp1 plays at least two temporally distinct roles in genome integrity.

These findings may provide insight into mammalian cancer development. P53 is one of the most important human tumor suppressors, mutated in more than 50% of human cancers (Muller & Vousden, 2014). Although p53 ensures genomic integrity as the key effector of the DDR (Levine et al., 2006; Kastenhuber & Lowe, 2017), it has been questioned whether this is the basis for tumor suppression. Point mutations that block p53 function in the acute DDR have little effect in blocking tumor growth, and conditional knock-out studies point to a critical window for tumor suppressor function that is subsequent to the resolution of DNA damage (Christophorou et al., 2005; Christophorou et al., 2006; Li et al., 2012; Kaiser & Attardi, 2018). In mammals, which seem to lack a clear Xrp1 ortholog, p53 is required for cell competition, potentially substituting functionally for the role of Xrp1 in *Drosophila* (Baker et al., 2019; Baker, 2020; Lacroix et al., 2020; Boutelle & Attardi, 2021; Pilley et al., 2021). Cell competition could provide a second role for mammalian p53 in genome integrity, contributing to tumor suppression, for example, if it were found to eliminate aneuploid cells, analogous to the way that Xrp1 contributes to both DDR and cell competition in *Drosophila*.

## RESULTS

### DNA damage-induced expression of Xrp1

Studies on irradiated *Drosophila* embryos and larval tissues indicate that DNA Damage-induced apoptosis proceeds in two distinct phases (Wells et al., 2006; Wichmann et al., 2006; McNamee & Brodsky, 2009; Ji et al., 2021a). An initial acute, p53-dependent DDR peaks in the first 4 hours (Figure 1A). This is followed by a later, p53-independent phase that is associated with cell competition (Figure 1A). Xrp1 mRNA is upregulated by p53 within 30 minutes of irradiation of *Drosophila* embryos (Brodsky et al., 2004; Akdemir et al., 2007), perhaps by direct binding of p53 within the Xrp1 locus (Kurtz et al., 2019). We explored Xrp1 mRNA and protein expression to confirm that Xrp1 is a p53 target in third-instar larval wing discs. We used a fly line in which the Xrp1 protein was expressed from the endogenous locus, containing a C-terminal HA tag (Blanco et al., 2020). Wild-type wing discs exhibited only a few Xrp1-HA-positive cells, possibly reflecting spontaneous DNA damage or other cellular stress (Figure 1B). Exposure to ionizing radiation (IR: 1500 Rads of γ-rays) induced strongly Xrp1-HA positive cells by 2 hours after IR, which further increased by 4 hours after IR (Figure 1C-D). The DNA damage-induced Xrp1-HA expression was non-uniform, with some cells showing higher expression levels than others. This heterogeneity was different from the more uniform Xrp1-HA expression seen in *Rp^+/-^* mutant wing discs (Figure 1E). A similar salt-and-pepper pattern of Xrp1 was observed with the Xrp1 antibody (Figure 1F, G). The specificity of Xrp1-HA expression was further confirmed by its reduction after RNAi for Xrp1 (Figure 1H).

**Figure 1:**
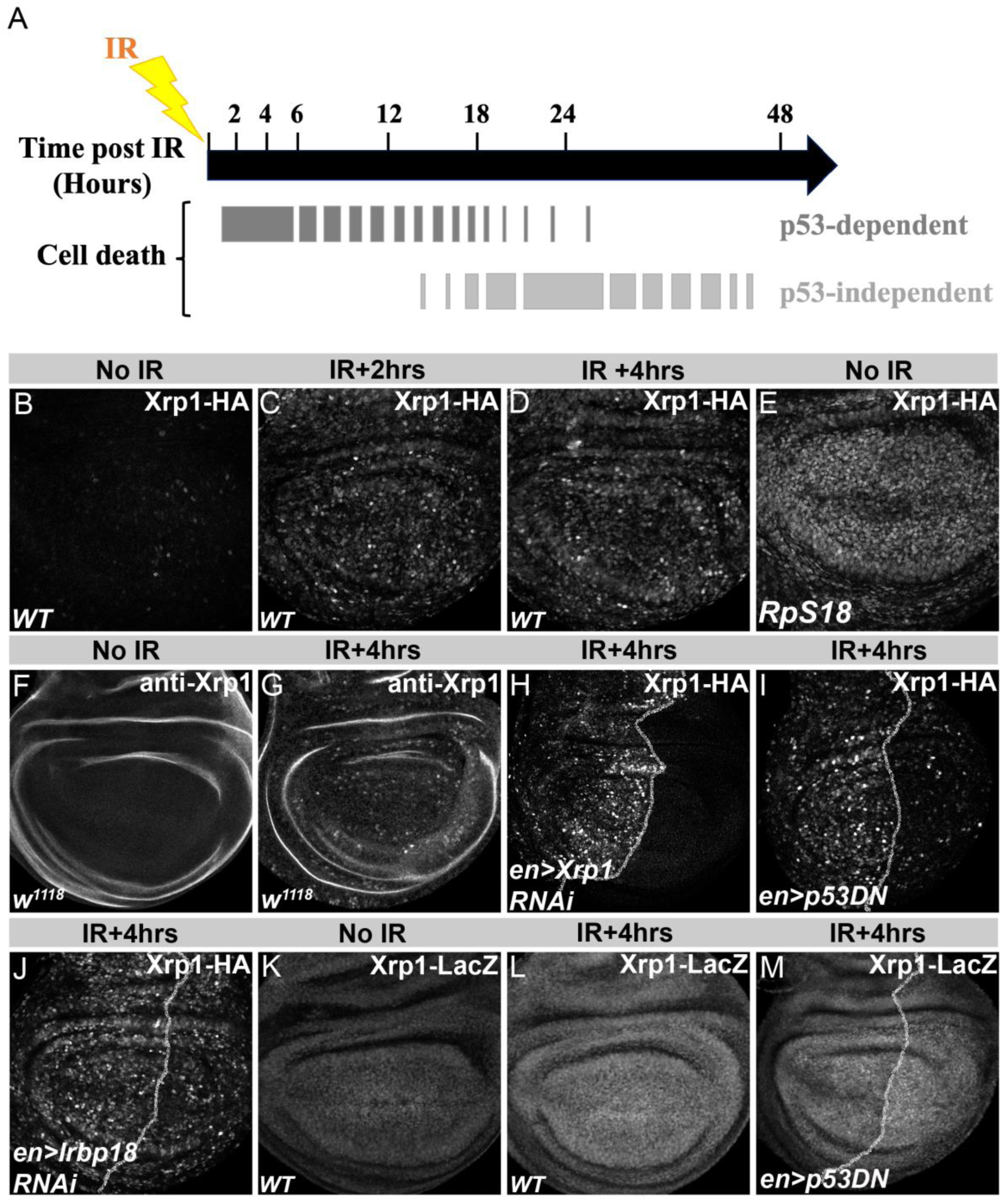
DNA damage induced Xrp1 expression. (A). Schematic of the approximate timeline of p53-dependent and p53-independent apoptotic responses following irradiation, based on previous work (Brodsky et al., 2004; Wells et al., 2006; McNamee & Brodsky, 2009; Ji et al., 2021b). (B). Xrp1 protein levels in the wild type III instar wing discs stained for HA, in 0 Rads (control) (C), 2 hours after 1500 Rads (D), 4 hours after 1500 Rads (E), and in *Rp18^+/-^* mutant (F). Larval wing discs stained with Xrp1 antibody, 0 Rads (control) (G), and 4 hours after 1500 Rads (H) (note: Xrp1 antibody shows higher background than Xrp1-HA). Xrp1-HA expression at 4 hours post-1500 Rads in the posterior compartment, compared relative to the anterior compartment, taken as an internal control, *en-Gal4* driving *UAS-Xrp1* RNAi (I), p53 dominant negative (*UAS-p53^H159N^*) (J), and *UAS-Irbp18* RNAi (K). III instar wing discs assayed for *Xrp1^02515^*-lacZ enhancer trap expression detected by anti-βgal antibody in 0 Rads (L), 4 hours after 1500 Rads (M), and in the wing disc expressing *UAS-p53^H159N^* under control of *en-Gal4,* 4 hours after 1500 Rads (N). N=7-8 discs were analyzed for every genotype, and each experiment was repeated at least 2-3 times. Genotypes: (B-D) *Xrp1-HA*, (E) *p{hs:FLP}; FRT42 RpS18 p{ubi:GFP},* (F-G) *w^1118^* (H) *en-Gal4::UAS-RFP, UAS-Xrp1 RNAi* (VDRC 107860)*; Xrp1-HA,* (I) *en-Gal4::UAS-RFP, UAS-p53DN^H159N^* (Bl. 8420)*; Xrp1-HA*, (J) *en-Gal4::UAS-RFP, UAS-Irbp18 RNAi; Xrp1-HA*, (K-L) *P{PZ}Xrp1^02515^*, (M) *en-Gal4::UAS-RFP,UAS-p53DN^H159N^* (Bl. 8420); *P{PZ}Xrp1^02515^*

We tested the requirement of p53 using a dominant-negative form encoded by the *UAS-p53^H159N^* transgene. We drove expression in posterior wing compartments so that Xrp1-HA expression levels could be compared to the anterior compartments of the same wing discs as a control. Dominant-negative p53 significantly reduced IR-induced Xrp1-HA (Figure 1I). Xrp1 in imaginal disc was also p53-dependent at the transcript level, with a ∼3x reduction in mRNA level seen after qRT-PCR from irradiated *p53^-/-^* wing discs (Figure 1- figure supplement 1A).

Interestingly, IR-induced Xrp1-HA protein expression was only slightly affected by Irbp18 knockdown (Figure 1J). This was different from Xrp1 expression in *Rp^+/-^* mutants, where Xrp1 autoregulation is important to regulate the level of Xrp1 expression (Baillon et al., 2018; Blanco et al., 2020). Consistent with autoregulation not contributing to IR-induced *Xrp1* transcription, 50% of the normal *Xrp1* mRNA level remained in the *Xrp1^m2-73^/Df* mutant background (Figure 1- figure supplement 1A). Thus, the *Xrp1^m2-73^* point mutant allele could still be transcribed in response to irradiation, even without a functional Xrp1 product.

An Xrp1-LacZ enhancer trap has also been used as a reporter previously. It is upregulated in *Rp^+/-^* mutant wing discs (Baillon et al., 2018; Lee et al., 2018a). Like Xrp1 protein and mRNA, Xrp1-LacZ was moderately elevated after irradiation (Figure 1K,L). Different from Xrp1-HA, Xrp1-LacZ expression depended on Xrp1 and Irbp18 function, both after irradiation and in the unirradiated control (Figure 1- figure supplement 1B-E). Unexpectedly, reducing p53 function slightly elevated the expression of Xrp1-LacZ after irradiation (Figure 1M). These differences might be explained if there is an autoregulatory component to Xrp1 transcription, particularly affecting Xrp1-LacZ, that does not contribute much to IR-induced Xrp1 mRNA or protein levels.

### Regulation of p53-dependent DDR target genes by Xrp1

In mammals, p53 regulates a complex transcriptional program of hundreds or even thousands of genes (Levine et al., 2006). Depending on the study, however, only 122-346 p53-dependent genes have a p53 consensus motif in the promoter or upstream region (Wei et al., 2006; Fischer, 2017). It is thought that p53 directly controls core transcription programs that, in turn, regulate further secondary, indirect targets (Andrysik et al., 2017). Xrp1 is a candidate for bringing secondary target genes under the indirect control of *Drosophila* p53. Consistent with this, when we compared 42 rapidly induced p53-target genes (RIPD) identified in the DDR (Brodsky et al., 2004; Akdemir et al., 2007), with Xrp1-dependent genes expressed in *Rp^+/-^* wing imaginal discs or discs overexpressing Xrp1 (Baillon et al., 2018; Lee et al., 2018b), we found 21 of 42 overlapped (Figure 2A). These genes were associated with Gene Ontology (GO) terms transcription factors, cell death, DNA repair, and redox response (Figure 2B).

**Figure 2:**
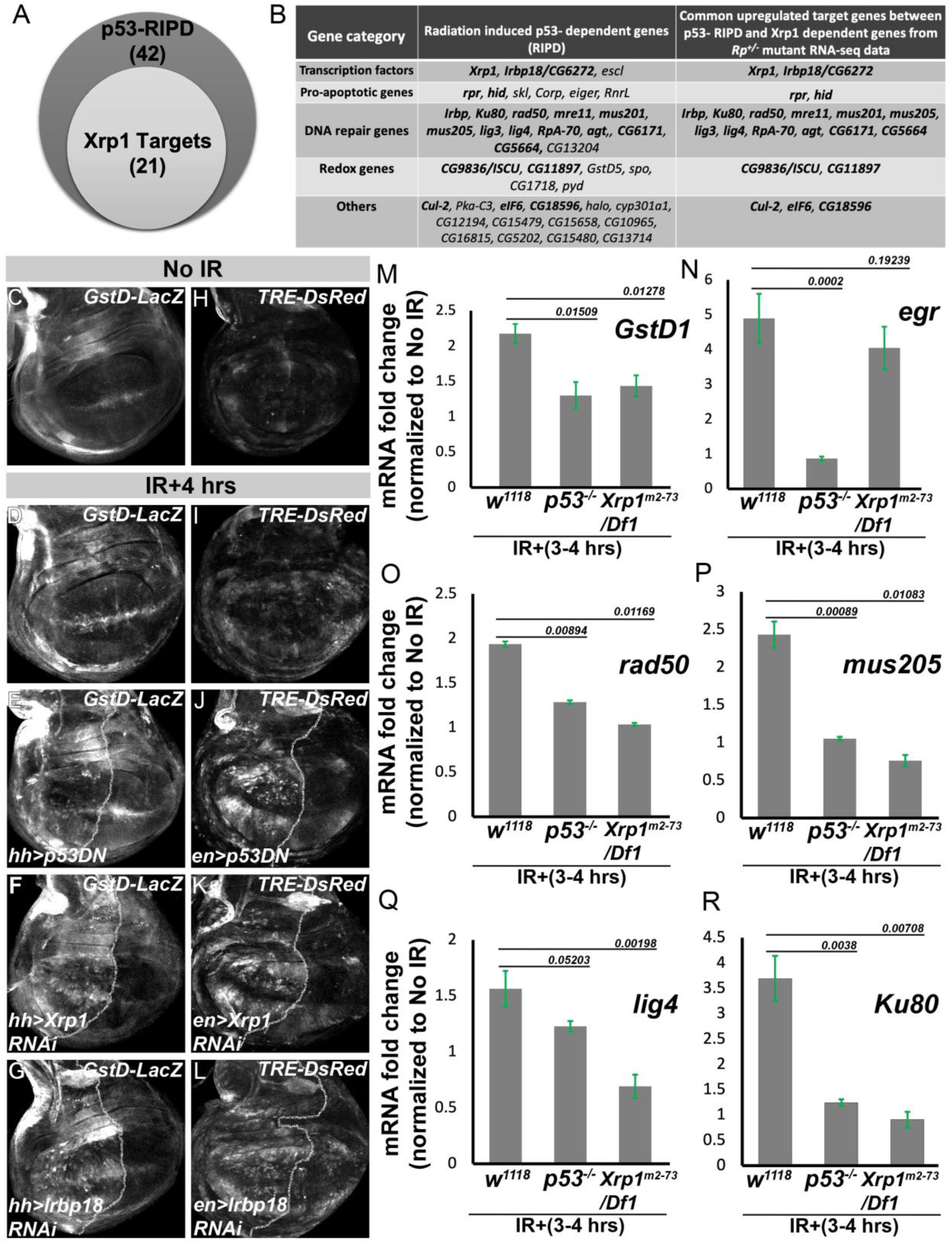
Xrp1 serves as the effector of p53 in DNA damage response. Vein diagram showing the overlap of p53 (Brodsky et al., 2004; Akdemir et al., 2007) and Xrp1 (Baillon et al., 2018; Lee et al., 2018b) target genes (A). The table shows the genes organized according to GO categories from radiation-induced p53-dependent (RIPD) and genes common between Xrp1- and p53-dependent transcription programs (highlighted in bold) (B). Expression of *GstD-*lacZ in III instar wing discs measured by staining for anti-βgal antibody, in the wild type (no IR) (C), 4 hours after 2000 Rads in wild type disc (D), *UAS-p53^H159N^* (E), *UAS-Xrp1 RNAi* (F), and *UAS-Irbp18* RNAi (G). Expression of *TRE-dsRed* in III instar wing discs measured by staining for anti-RFP antibody, in the wild type (no IR) (H), 4 hours after 1500 Rads in wild type disc (I), *UAS-p53^H159N^* (J). *UAS-Xrp1 RNAi* (K) and *UAS-Irbp18 RNAi* (L). qRT-PCR analysis for mRNA expression levels from III instar wing discs of *w^1118^*, *p53^[5A-1-4]^* and *Xrp1^m2-73^/Df1* collected 3-4 hours after irradiation (1500 RADs) and normalized to untreated (0 Rads) sample, for the following genes *GstD1* (M), *egr* (N), *rad50* (O), *mus205* (P), *lig4* (Q) and *Ku80* (R). N=7-8 discs were analyzed for every genotype, and each experiment was repeated at least 2-3 times. Data is plotted as mean± SEM, a two-tailed student t-test for independent means, was performed for statistical significance. Genotype: (C & D) *GstD-LacZ,* (E) *GstD-LacZ, UAS-p53^H159N^* (Bl. 8420)*; hh-Gal4::UAS-GFP,* (F) *GstD-LacZ, UAS-Xrp1 RNAi* (VDRC 107860)*; hh-Gal4::UAS-GFP,* (G) *GstD-LacZ, UAS-Irbp18 RNAi; hh-Gal4::UAS-GFP*, (H and I) *TRE-dsRed,* (J) *TRE-dsRed::en-Gal4, UAS-p53^H159N^* (Bl. 8420), (K) *TRE-dsRed::en-Gal4, UAS-Xrp1 RNAi* (VDRC 107860), (L) *TRE-dsRed::en-Gal4, UAS-Irbp18 RNAi*, (M-R) *w^1118^*, *yw; p53^[5A-1-4]^,* and *Xrp1^m2-73^/Df1*

To test if Xrp1 was needed to express p53-dependent genes in response to IR, we employed available reporter lines and also confirmed gene expression by qRT-PCR. *GstD-lacZ* is a redox response reporter that is induced in response to oxidative stress (Sykiotis & Bohmann, 2008; Brown et al., 2021) and is elevated by Xrp1 in *Rp^+/-^* wing discs (Kucinski et al., 2017; Ji et al., 2019; Kiparaki et al., 2022). *GstD-lacZ* was expressed in response to IR, dependent on p53 (Figure 2C-D). We found that Xrp1 and Irbp18 were needed (Figure 2F-G). This was in accordance with Xrp1/Irbp18 binding sites being present in the *GstD*-lacZ reporter (Brown et al., 2021). To confirm that antioxidant genes themselves were affected like the *GstD-lacZ* reporter was, we measured *GstD1* mRNA levels in wing discs. Induction of *GstD1* mRNA by IR indeed depended on both p53 and Xrp1 (Figure 2M).

Another stress pathway activated in DDR is the Jun N-terminal Kinase (JNK) pathway (Picco & Pages, 2013). JNK activity is also Xrp1-dependent in *Rp^+/-^* wing discs (Lee et al., 2018b; Ji et al., 2019). We found the JNK reporter *TRE-dsRed* was induced by IR in a p53-dependent manner (Figure 2H-J), and this was Xrp1/Irbp18 dependent (Figure 2K & L). The TNF-alpha ligand *eiger (egr)* has been proposed to activate JNK signaling in the DDR (Sanchez et al., 2019). Surprisingly, however, *egr* transcription was p53-dependent but independent of Xrp1, which questions whether it is solely responsible for JNK signaling in the DDR (Fig. 2N).

All except one of the p53-dependent DNA repair genes induced in DDR were also potential Xrp1 targets (Figure 2B). We measured wing disc mRNA levels of representative genes from the major DNA repair pathways. These were the homologous recombination pathway gene *rad50*, the translesion polymerase DNA polymerase zeta subunit 1, *mus205*, and the Non-Homologous End Joining (NHEJ) pathway genes DNA ligase 4 (*lig4*) and *Ku80*. We found that IR-dependent expression of all these genes depended on Xrp1 at least as strongly as upon p53 (Figure 2O-R).

A well-known aspect of DDR is the apoptotic death of severely damaged cells, dependent in *Drosophila* on p53-dependent induction of the pro-apoptotic genes *rpr* and *hid* (Brodsky et al., 2004; Dekanty et al., 2015). To explore the potential role of Xrp1, we examined reporter lines for *rpr* and *hid*, which are expressed in response to IR and are widely used as p53 reporters (Brodsky et al., 2004; Tanaka-Matakatsu et al., 2009; Fan et al., 2010). Two reporter lines for *hid* are the *hid5’F-wt-EGFP* reporter, which contains ∼2 kb spanning the *hid* promoter from +16 to −2210bp relative to the transcription start site, and the *hid^20-10^-lacZ* reporter, which includes a distinct 10 kb sequence from the *hid* upstream region (Tanaka-Matakatsu et al., 2009; Fan et al., 2010) (Figure 3A). As expected, *hid5’F-wt-EGFP* was highly induced after IR in a p53-dependent manner (Figure 3B-D). Both *Xrp1* RNAi and *Irbp18* RNAi reduced the *hid5’F-wt-EGFP* expression to the same extent as p53 dominant-negative did (Figure 3E-F). The *hid^20-10^-lacZ* reporter was also induced by IR and was dependent upon p53 (Figure 3G-I). Both *Xrp1* and *Irbp18* RNAi reduced IR-dependent *hid^20-10^-lacZ* expression, although not so much as p53 dominant-negative (Figure 3J-K). We assessed *rpr* expression using the *rpr^150^-lacZ* reporter, containing a 150bp sequence including a putative p53 binding site ∼4.8kb upstream of the *rpr* gene (Brodsky et al., 2000). In controls, *rpr^150^-lacZ* reporter exhibits uniform expression in most of the wing disc, on which is superimposed a region of elevated expression bordering the posterior wing pouch and notum region (Alpar et al., 2018; Li et al., 2020) (Figure 3L). It is now appreciated that many DDR genes have non-uniform expression patterns in the wing disc (Cruz et al., 2025). Because IR enhanced all aspects of *rpr^150^-lacZ* expression (Figure 3L, M, and Figure 3- figure supplement 1A), we measured the average *rpr^150^-lacZ* expression level in anterior and posterior compartments and computed the ratio of posterior-to-anterior expression level as a readout of genetic perturbations applied to the posterior compartments (Figure 3- figure supplement 1B). The increase in *rpr^150^-lacZ* following IR was p53-dependent, as expected, but unaffected by RNAi for either Xrp1 or Irbp18 (Figure 3N-P- figure supplement 1B).

**Figure 3:**
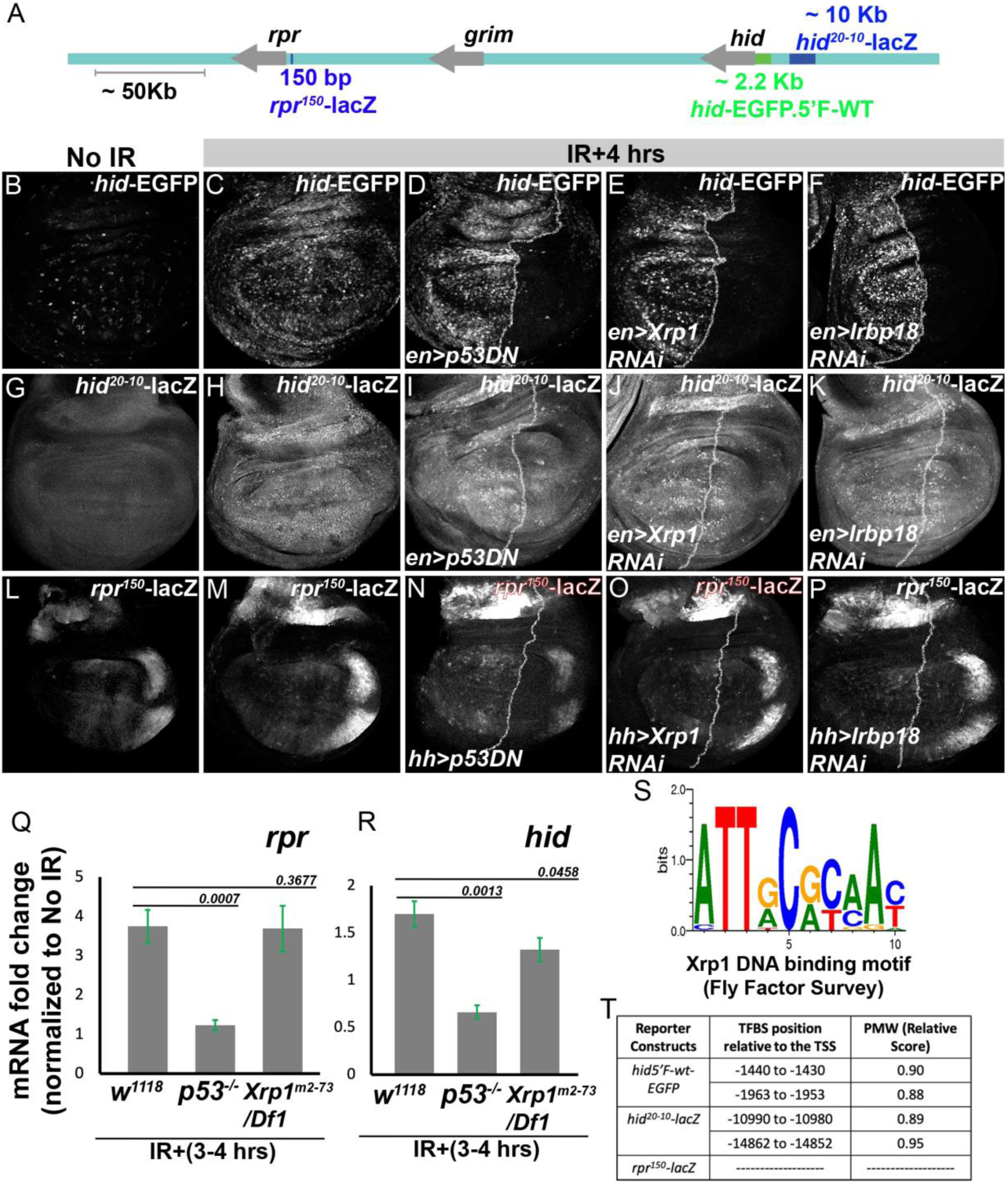
Xrp1 controls the expression of the regulatory sequences of the pro-apoptotic gene *hid*. Schematic of H99 locus with approximate positions of the *rpr, hid,* and *grm* and the reporter lines used for the analysis (A). Expression of *hid5’F-wt-EGFP* measured by staining for anti-GFP antibody, in the wild type (no IR) (B) and 4 hours after 2000 Rads in the wild type wing disc (C), *UAS-p53^H159N^* (D), *UAS-Xrp1 RNAi* (E), and *UAS-Irbp18* RNAi (F). *hid^20-10^-lacZ* expression measured by anti-βgal staining in the wild type (no IR) (G) and 4 hours after 2000 Rads in wild type wing disc (H), *UAS-p53^H159N^* (I), *UAS-Xrp1* (J), and *UAS-Irbp18* RNAi (K). *rpr^150^-lacZ* expression measured by anti-βgal staining, in the wild type (no IR) (L) and 4 hours after 2000 Rads in the wild type wing disc (M). *UAS-p53^H159N^* (N), *UAS-Xrp1* (O), and *UAS-Irbp18* RNAi (P). qRT-PCR analysis for mRNA expression levels from III instar wing discs of *w^1118^*, *p53^[5A-1-4]^*, and *Xrp1^m2-73^/Df1* collected 3-4 hours after irradiation (1500 RADs) and normalized to untreated (0 Rads) sample, for the following genes, *rpr* (Q), and *hid* (R). Putative DNA binding motif of Xrp1/Irbp18 heterodimer, adapted from Fly Factor Survey (S). Position of Xrp1/Irbp18 DNA motif in the upstream regulatory region of *hid* relative to the TSS (T). N=5-8 discs were analyzed for every genotype; data was plotted as mean± SEM, and a two-tailed student t-test for independent means was performed for statistical significance. Genotype: (B & C) *hid5’F-wt-EGFP*, (D) *hid5’F-wt-EGFP; en-Gal4::UAS-RFP, UAS-p53^H159N^* (Bl. 8420), (E) *hid5’F-wt-EGFP; en-Gal4::UAS-RFP, UAS-Xrp1 RNAi* (VDRC 107860), (F) *hid5’F-wt-EGFP; en-Gal4::UAS-RFP, UAS-Irbp18 RNAi*, (G & H) *yw*; *hid^20-10^-lacZ*, (I) *en-Gal4::UAS-RFP, UAS-p53^H159N^* (Bl. 8420)*; hid^20-10^-lacZ*, (J) *en-Gal4::UAS-RFP, UAS-Xrp1 RNAi* (VDRC 107860)*; hid^20-10^-lacZ*, (K) *en-Gal4::UAS-RFP, UAS-Irbp18 RNAi; hid^20-10^-lacZ*, (L & M) *rpr^150^*-lacZ, (N) *rpr^150^-lacZ, UAS-p53^H159N^* (Bl. 8420)*; hh-Gal4::UAS-GFP*, (O) *rpr^150^-lacZ, UAS-Xrp1 RNAi* (VDRC 107860)*; hh-Gal4::UAS-GFP,* (P) *rpr^150^-lacZ, UAS-Irbp18 RNAi; hh-Gal4::UAS-GFP*, (Q-R) *w^1118^*, *yw; p53^[5A-1-4]^,* and *Xrp1^m2-73^/Df1*

To confirm that these studies of *hid* and *rpr* reporters were indeed reflective of the expression of the genes themselves, mRNA levels were measured in wing discs after IR. Consistent with the reporter studies, induction of *hid* mRNA by irradiation was completely dependent on p53 and partially dependent on Xrp1 (Figure 3Q). Induction of *rpr* mRNA expression was completely dependent on p53 but was unaffected by the loss of Xrp1 (Figure 3R).

Taken together, these results show that genes identified as p53 targets in the *Drosophila* DDR include genes that are, in fact, directly induced by Xrp1. Only two out of nine p53 target genes examined, *egr* and *rpr*, were induced independently of Xrp1, although *hid* induction has both Xrp1-dependent and -independent components. Analysis of p53 and Xrp1 binding motifs supported this conclusion. Sequences matching the p53 binding consensus have been located within both the *rpr^150^-lacZ* reporter and the *hid5’F-wt-EGFP* reporter (Brodsky et al., 2000; Wichmann et al., 2010). Consensus binding sites for the Xrp1/Irbp18 heterodimer have been determined by bacterial one-hybrid studies (Fly Factor Survey) (Zhu et al., 2011) and from ChIP-seq after Xrp1 overexpression in wing disc (Baillon et al., 2018). Such sequences are footprinted by Xrp1/Irbp18 in the P element terminal repeat (Francis et al., 2016) and are sufficient to confer Xrp1/Irbp18-dependent transcription on luciferase reporters (Kiparaki et al., 2022). Using an *in-silico* motif finder (INSECT) (Rohr et al., 2013), we found two Xrp1/Irbp18 binding motifs in the *hid5’F-wt-EGFP* reporter sequence and two in the *hid^20-10^-lacZ* reporter sequence (Figure 3T), but we found none in the *rpr^150^-lacZ* reporter that was expressed independently of Xrp1 and Irbp18 (Figure 3T). We conclude that, while *rpr* could be a direct target of p53 in the DDR, *hid* expression is partially dependent on Xrp1 induction downstream of p53, and the *hid5’F-wt-EGFP* reporter entirely so.

### Xrp1 is sufficient for the expression of p53-dependent DDR target genes

It was possible that the only Xrp1 requirement in the DDR might be to activate p53 to create a positive feedback loop. We used the *rpr^150^-lacZ* reporter, which is activated by p53 but not Xrp1, to test how p53 activity was affected by Xrp1 overexpression. Xrp1 overexpression in the wing pouch resulted in a huge amount of cell death, as expected (Figure 4- figure supplement 1A-B), but no discernible increase in *rpr^150^-lacZ* levels (Figure 4- figure supplement 1C-D). Similar results were obtained by expressing Xrp1 in the eye discs (Fig. 4A-B). Thus, Xrp1 did not cause p53 activation and is likely to act downstream of p53.

**Figure 4:**
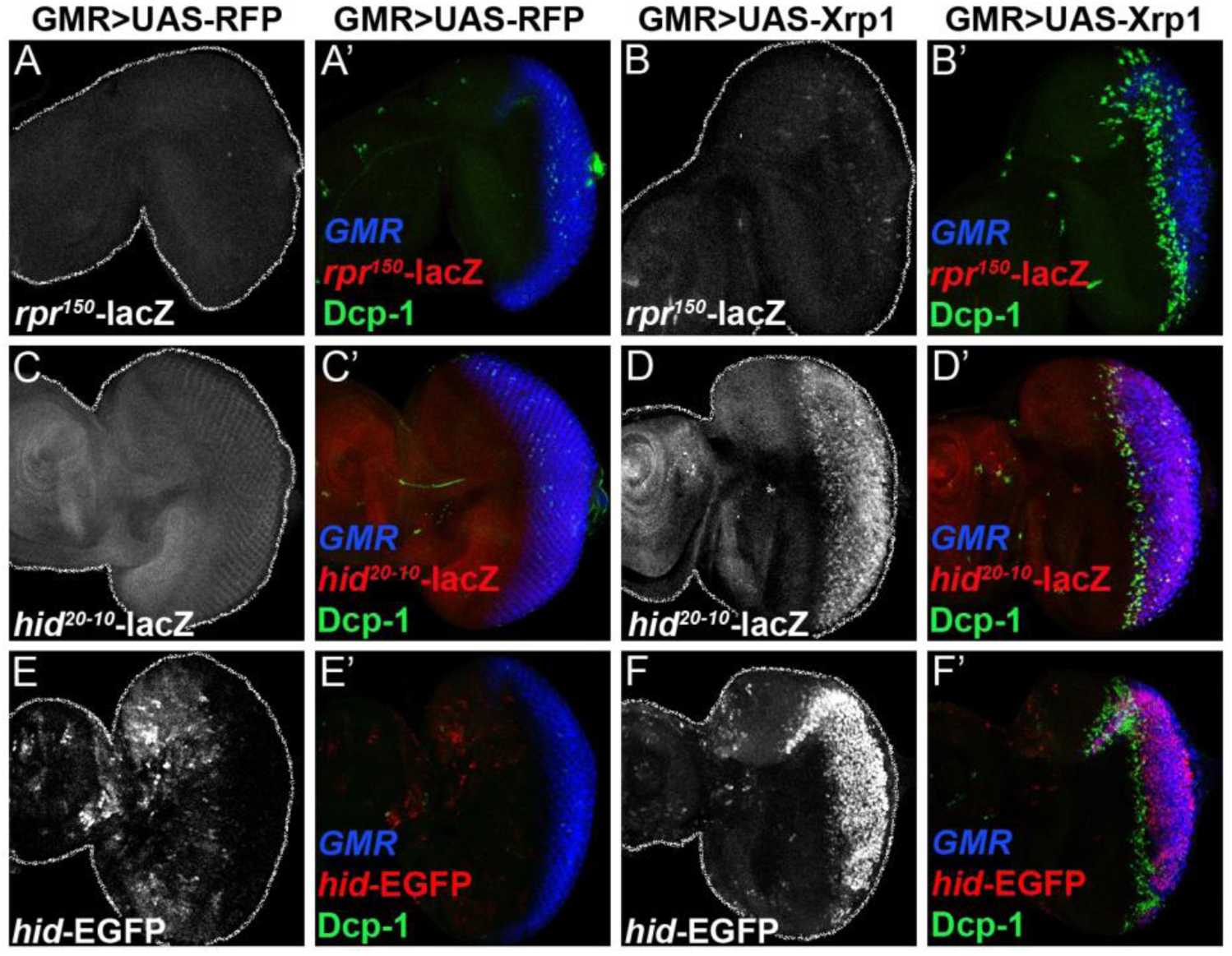
Xrp1 is sufficient for the expression of p53-dependent DDR target genes. *rpr^150^-lacZ* reporter expression measured by anti-βgal (red) and cell death by anti-Dcp-1 (green) in III instar eye imaginal disc expressing UAS-RFP (A) and UAS-Xrp1 long isoform (B), under the control of *GMR-Gal4* (blue). *hid^20-10^-lacZ* reporter expression measured by anti-βgal (red) and cell death by anti-Dcp-1 (green) in III instar eye imaginal disc expressing UAS-RFP (C) and UAS-Xrp1 long isoform (D) under the control of *GMR-Gal4* (blue). *hid5’F-wt-EGFP* reporter expression measured by anti-GFP (red) and cell death by anti-Dcp-1 (green) in III instar eye imaginal disc expressing UAS-RFP (E) and UAS-Xrp1 long isoform (F) under the control of *GMR-Gal4* (blue). N=7-8 discs were analyzed for every genotype; each experiment was repeated at least 2 times. All the control and experimental genotypes shown in this figure were kept at 18°C to lessen the effect of Xrp1 overexpression. Genotype: (A-A’) *GMR-Gal4::UAS-RFP, rpr^150^-lacZ,* (B-B’) *GMR-Gal4::UAS-RFP::UAS-Xrp1, rpr^150^-lacZ,* (C-C’) *GMR-Gal4::UAS-RFP; hid^20-10^-lacZ,* (D-D’) *GMR-Gal4::UAS-RFP::UAS-Xrp1; hid^20-10^-lacZ,* (E-E’) *hid5’F-wt-EGFP; GMR-Gal4::UAS-RFP*, (F-F’) *hid5’F-wt-EGFP; GMR-Gal4::UAS-RFP, UAS-Xrp1*

Next, we tested if Xrp1 was sufficient for the expression of target genes. Xrp1 overexpression in the eye disc led to a robust upregulation of *hid^20-10^-lacZ* (Figure 4C and D) and *hid5’F-wt-EGFP* (Figure 4E and F). Xrp1 also regulated JNK and *GstD* after DNA damage, and Xrp1 overexpression was sufficient to activate the reporters for these pathways, *TRE-dsRED* and *GstD-lacZ* (Figure 4- figure supplement 1E and F & G and H). These results show that Xrp1 functions downstream of p53 and is sufficient for expressing a subset of p53-dependent DDR genes.

### Xrp1 contributes to DNA damage-induced cell death

To see whether the regulation of cell death genes by Xrp1 was functionally significant, the *p53*-dependent apoptosis that occurs in response to irradiation was examined. Cell death begins within 30 minutes after IR treatment and is entirely dependent on p53 for at least 4-5 hours (Wichmann et al., 2006; McNamee & Brodsky, 2009). Accordingly, p53 dominant-negative expression in posterior compartments substantially reduced the IR-induced cell death, leaving only a few cells positive for the activated Dcp-1 caspase (Figure 5A, B, D). Xrp1 RNAi (two independent lines) and Irbp18 RNAi both reduced cell death in the posterior compartment (Figure 5C, D) to a lesser extent than p53 dominant-negative did.

**Figure 5:**
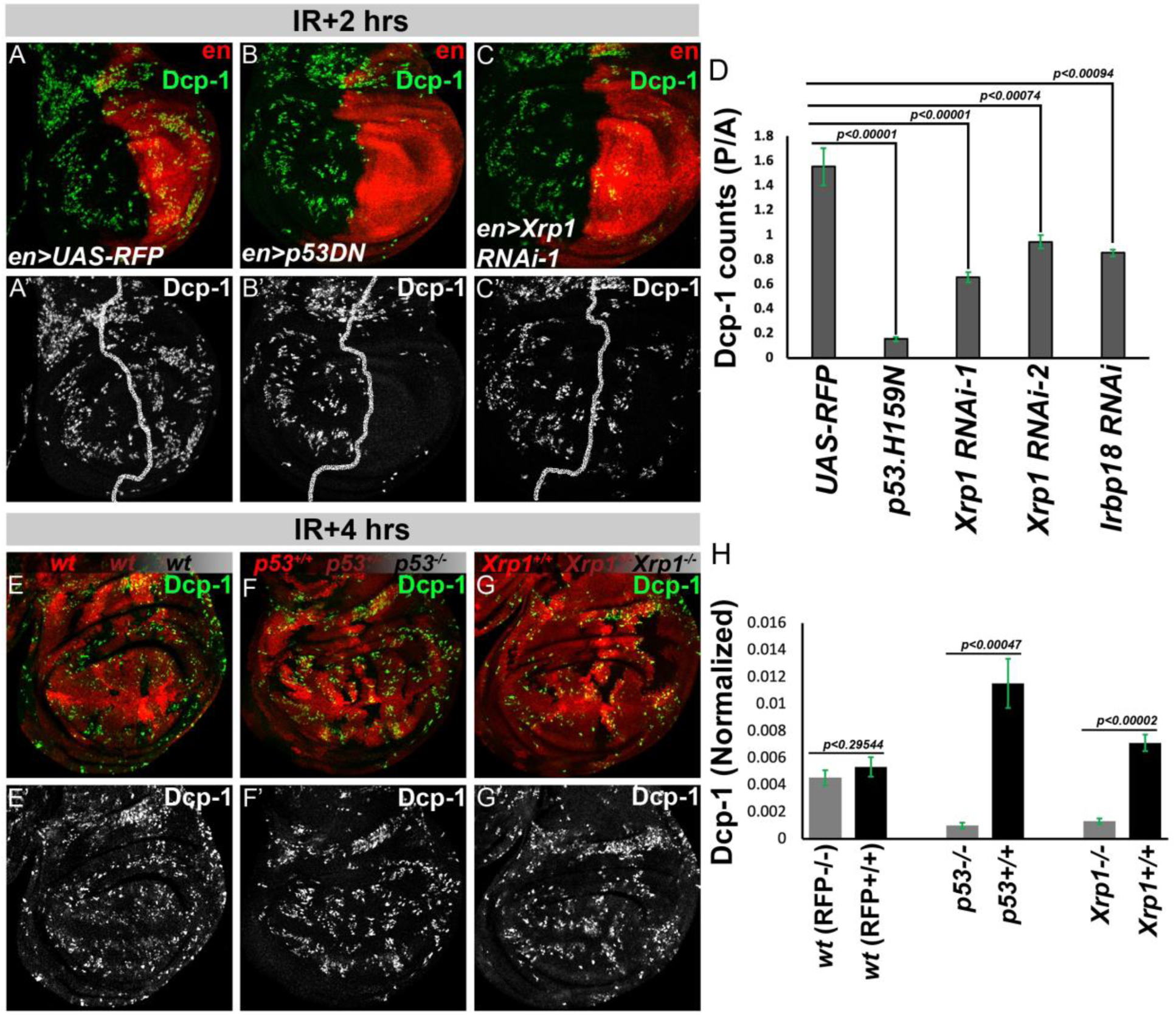
Xrp1 is partly required for DNA damage-induced cell death. *Drosophila* III instar wing imaginal discs dissected at 2 hours after 1500 Rads, cell death is measured by Dcp-1 (green and grayscale) in different genetic perturbations using en-Gal4 (posterior compartment (red)), UAS-RFP (A), *UAS-p53^H159N^* (B), and *UAS-Xrp1* RNAi (C). Dcp-1 positive cells were quantified and plotted as the ratio of Dcp-1 positive cells in posterior (P) to anterior (A) compartments (D). Wing discs were dissected 4 hours after 1500 RADs and stained for Dcp-1 positive cells in *FRT82* (E), *FRT82p53^[5A-1-4]^* (F), and *FRT82Xrp1^m2-73^* (G). (H) Dcp-1 positive cells were quantified in the clone (RFP-/-) and in the sister clone (RFP+/+). A 4-hour timepoint was taken to have a uniform distribution of Dcp-1 positive cells. Statistics: (A-D) N=15 for each genotype. (E-H) N=8 for each genotype. Data is plotted as mean± SEM, a two-tailed student t-test for independent means for (A-D) and dependent means for (E-H), was performed for statistical significance. Genotype: (A) *en-Gal4::UAS-RFP,* (B) *en-Gal4::UAS-RFP, UAS-p53DN^H159N^* (Bl. 8420), (C) *en-Gal4::UAS-RFP, UAS-Xrp1 RNAi* (Bl. 4521), (E) *p{hs:FLP}; FRT82-RFP/FRT82* (30 min hs, 48 hr A.E.L), (F) *p{hs:FLP}; FRT82-RFP/FRT82-p53^[5A-1-4]^* (30 min hs, 48 hr A.E.L), (G) *p{hs:FLP}; FRT82-RFP/FRT82-Xrp1^m2-73^* (30 min hs, 48 hr A.E.L)

We also examined the cell autonomy of p53 and Xrp1 function using clones of mutant cells. Control clones did not affect IR-induced cell death (Figure 5E, H). Wild-type and *p53^+/-^* heterozygous cells exhibited similar cell death levels after irradiation, but cell death was absent from *p53^-/-^* null clones (Figure 5F, H). There was little cell death observed in *Xrp1^+/-^* (heterozygous) and *Xrp1^-/-^* (null) tissue as opposed to the high amount of cell death observed in *Xrp1^+/+^* (wild type twin spot) (Figure 5G, H) (effects of Xrp1 mutation on *Rp* mutant cells are also dominant through haploinsufficiency) (Lee et al., 2018b). Taken together, these results show that Xrp1 contributes to p53-mediated cell death in response to irradiation, although this is partial because p53 also regulates cell death genes directly, such as *rpr* (Figure 3R).

We also tested the contribution of Xrp1 to the effects of p53 overexpression. Over-expression of p53 in the developing eye causes a rough eye phenotype in adults (Figure 5- figure supplement 1A) (Jin et al., 2000; Ollmann et al., 2000). Xrp1 knockdown very slightly reduced the rough eye phenotype, unlike dominant-negative p53, which rescued completely (Figure 5- figure supplement 1B, C). p53 overexpression in wing discs resulted in massive cell death that was reduced only weakly by Xrp1 knockdown or mutant genotypes (Figure 5- figure supplement 1D-H). In accordance with this partial dependence on Xrp1, p53 overexpression led to rather little Xrp1 protein expression (Figure 5- figure supplement 1I). These differences suggest that the effects of p53 overexpression differ from endogenous p53 activity in the physiological DDR, which depends on Xrp1 to a considerable degree.

### Xrp1-dependent competitive cell death limits genomic instability

The DDR is followed by a phase of p53-independent cell death, peaking around 24h after irradiation, reflecting the loss of aneuploid cells (Wichmann et al., 2006; McNamee & Brodsky, 2009). We previously showed that the majority of this p53-independent cell death depends on RpS12 and is associated with the expression of Xrp1, consistent with cell competition on the basis of altered *Rp* gene dose (Ji et al., 2021b). Here, we confirmed that most of the p53-independent cell death depends on Xrp1 (Figure 6A-F). At 24 hours after irradiation, Xrp1-HA expression was still observed and was totally suppressed upon Xrp1 knockdown (Figure 6- figure supplement 1A-C). Irbp18 knockdown reduced Xrp1-HA expression to some extent in accordance with the autoregulatory role of Irbp18/Xrp1 in cell competition (Figure 6- figure supplement 1D, G). It was already known that Xrp1 expression at this stage did not require p53 (Ji et al., 2021a); surprisingly, Xrp1-HA expression was actually increased when p53 was inhibited (Figure 6- figure supplement 1E, G). This increase was reduced in the *rpS12* mutant (Figure 6- figure supplement 1F, G). Importantly, in addition to not affecting cell death at 2 hours after IR (Figure 6- figure supplement 1H, I) (Ji et al., 2021a), RpS12 also does not affect Xrp1-HA expression at 2 hours after IR (Figure 6- figure supplement 1J-L). Thus, Xrp1 expression and function in the acute DDR depended on p53, not RpS12. In contrast, the Xrp1 expression that occured 24 hours after irradiation depended at least in part on RpS12, as did the enhanced expression that occured in the absence of p53.

**Figure 6:**
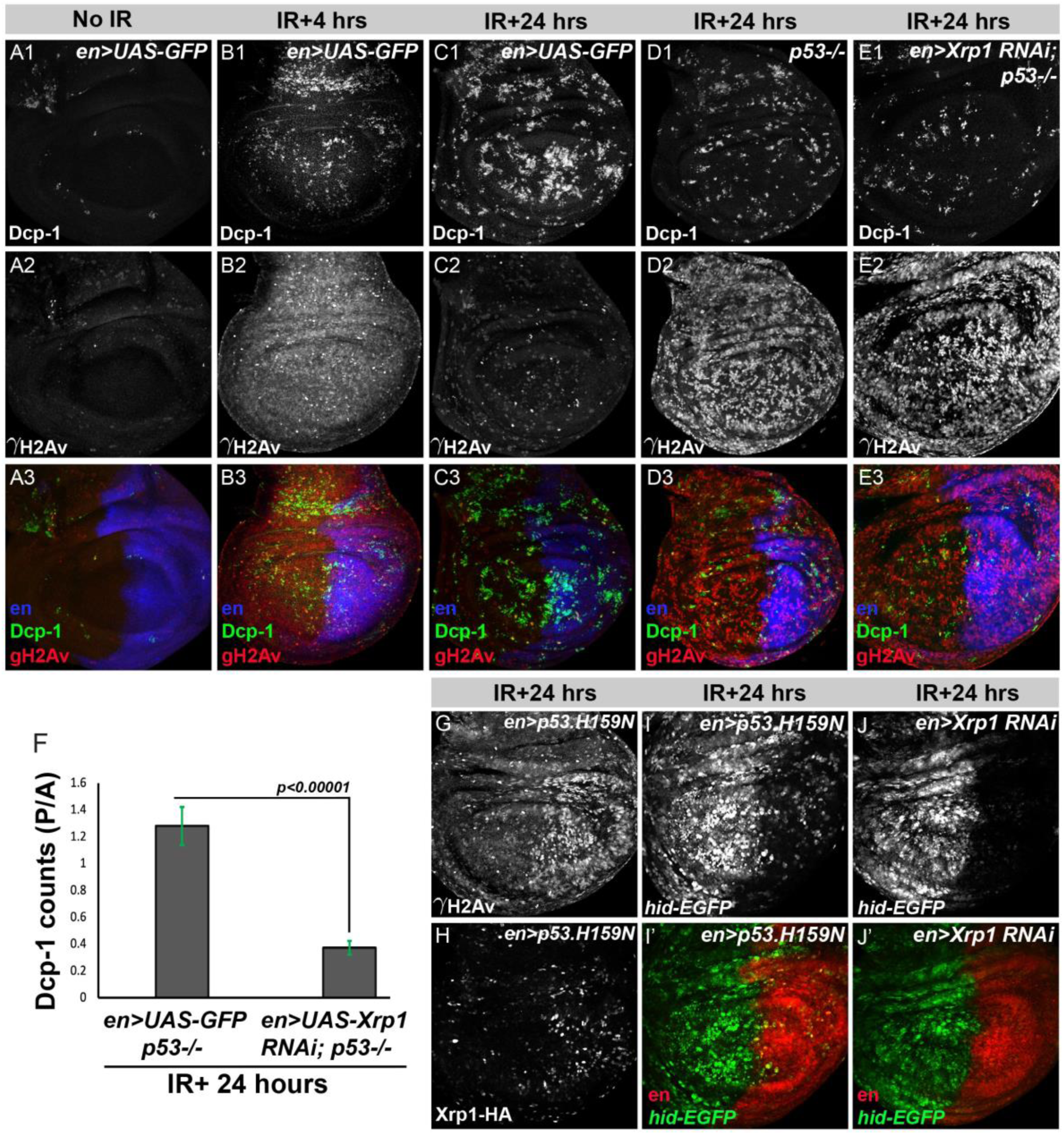
Xrp1 limits genomic instabilities by removing aneuploid cells in a p53-independent manner. (A-B) III instar wing disc from larvae stained for cell death (Dcp-1, green), DNA damage signaling (γH2Av, red), control (*en-Gal4, UAS-GFP*) (A), and 4 hours after 1500 Rads of γ-rays treatment (B). (C-E) III instar wing disc from larvae 24 hours after 1500 Rads of γ-rays treatment, stained for cell death (Dcp-1, green), DNA damage signaling (γH2Av, red) in control (*en-Gal4, UAS-GFP*) (C), *p53^-/-^* mutant (D), and Xrp1 k/d in *p53^-/-^* mutant background (E). (F) Quantification of Dcp-1 positive cells anterior versus posterior compartments in *p53^-/-^* mutant and Xrp1 k/d in *p53^-/-^* mutant background. (G) γH2Av and (H) Xrp1-HA levels 24 hours after irradiation in the posterior compartment of the wing disc expressing *p53^H159N^* under the control of engrailed Gal4*. hid5’F-wt-EGFP* expression in *UAS-p53^H159N^* (I) and Xrp1 RNAi K/D (J) 24 hours after irradiation. N=15 for (A-C), and N=8, for (E & F). Data is plotted as mean± SEM, student t-test, two-tailed for the independent sample, was performed for statistical significance. Genotype: (A-B) *en-Gal4::UAS-GFP*, (C) *en-Gal4::UAS-GFP; FRT82-RFP/FRT82-p53^[5A-1-4]^*, (D) *en-Gal4::UAS-GFP; FRT82-RFP/FRT82-p53^[5A-1-4]^*, (E) *en-Gal4::UAS-GFP, UAS-Xrp1 RNAi* (VDRC 107860)*; FRT82-RFP/FRT82-p53^[5A-1-4]^,* (G) *en-Gal4::UAS-RFP; UAS-p53DN^H159N^* (Bl. 8420)*; Xrp1-HA*, (H) *en-Gal4::UAS-RFP; UAS-p53DN^H159N^* (Bl. 8420)*; Xrp1-HA*, (I) *hid5’F-wt-EGFP; en-Gal4::UAS-RFP, UAS-p53^H159N^* (Bl. 8420), and (J) *hid5’F-wt-EGFP; en-Gal4::UAS-RFP, UAS-Xrp1 RNAi* (VDRC 107860)

To explore the impacts of p53 on genome integrity and Xrp1 activity further, we looked at IR-induced DNA damage using antibodies against γH2Av, which marks the double-strand breaks (Lake et al., 2013; Khan et al., 2017). Consistent with previous reports, most of the acute DNA damage visible 4h after irradiation was repaired by 24 hours (Figure 6A-C) (Wells et al., 2006; McNamee & Brodsky, 2009). Like Xrp1-HA expression, γH2Av levels 24 hours after irradiation were increased in the absence of p53, indicating persistent unrepaired DNA damage (Figure 6D). Dual Xrp1 knockdown and p53 inhibition led to an even further increase (Figure 6E). These data indicated that p53 was required for the repair (or removal) of irradiation-damaged DNA, and that some cells where this damage persists were then removed by p53-independent Xrp1 activity, i.e., cell competition. Consistent with this conclusion, double labeling showed that some of the γH2Av-positive cells that persisted when p53 was inhibited also expressed Xrp1-HA (Figure 6G-H). In support of the p53-independent function of Xrp1 at 24 hours post-IR, the expression of *hid5’F-wt-EGFP* was totally dependent on Xrp1 at this stage, but not all this expression depended on p53 (Figure 6J, I). However, we noticed that a fraction of *hid5’F-wt-EGFP* expression at 24 hours post-IR was still p53-depedent but to a much lesser extent that at 4 hours (Figure 3).

To get a better idea of when cell competition begins, wing discs were examined 16 hours post-IR. We found that Xrp1-HA expression was largely independent of p53 by 16 hours (Figure 6- figure supplement 2A-C) and, in fact, similar to 24 hours, Xrp1 was elevated in the absence of p53. However, similar to 4 hours after IR, Xrp1 knockdown completed suppressed Xrp1-HA expression at 16 hours post-IR but Irpb18 did not (Figure 6- figure supplement 2D, E). Cell death occurring 16 hours after irradiation was substantially p53-dependent, but p53-independent cell death was also detected (Figure 6- figure supplement 2F, G & I), in contrast to the acute DDR phase, where cell death is almost completely p53 dependent (Figure 5). Cell death 16 hours following irradiation was reduced by Xrp1 and Irbp18 depletion, but not eliminated (Figure 6- figure supplement 2F, H & I). These results suggest that p53-independent Xrp1 expression and function in cell competition are emerging by 16 hours and peaking by 24 hours after irradiation, perhaps indicating the timeline of the appearance of aneuploid cells.

## DISCUSSION

Xrp1 is known to contribute to genome integrity because flies lacking Xrp1 exhibit increased loss-of-heterozygosity following irradiation (Akdemir et al., 2007; Ji et al., 2021b). Here, we affirm that Xrp1 contributes to genome maintenance at two levels (Figure 7). One is as a component of the acute DDR, mediating part of the p53 transcriptional response and p53-dependent apoptosis; the other role occurs later, through cell competition to eliminate aneuploid and segmentally aneuploid cells that have acquired altered Rp gene dose as a result of DNA damage and its subsequent repair (Figure 7A). Xrp1 expression is induced by p53 in the DDR and by RpS12 during cell competition. Defects in the DDR, eg, when p53 is inhibited, increase the aneuploid cell burden and the resulting need for cell competition (Figure 7B,C). These two processes are distinguished in *Drosophila* by the distinct roles of p53 and of RpS12 in Xrp1 induction in each case. In mammals, however, both the DDR and cell competition depend on p53 and are both potential contributors to p53-mediated tumor suppression.

**Figure 7:**
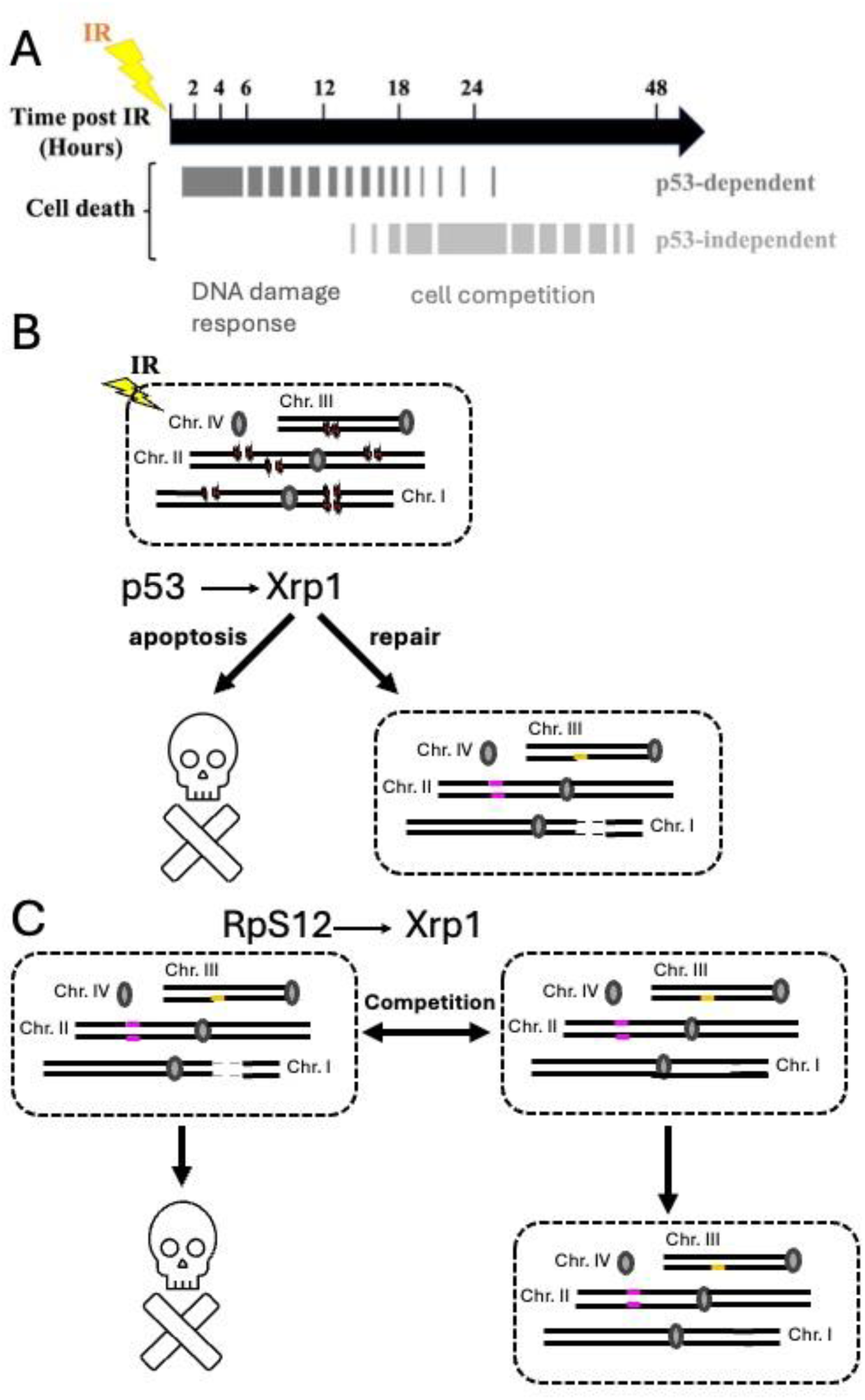
Model. (A) Our work reinforces the conclusion that an acute, p53-dependent apoptotic response to irradiation represents the DDR, while the later, p53-independent phase is associated with elimination of aneuploid cells by cell competition. The p53-independent phase depends more completely on Xrp1, whose expression at this stage is RpS12-dependent. The cell competition response is enhanced when p53 is absent, resulting in prolonged DNA damage and presumably an increase in aneuploidy. (B) The acute, p53-dependent DDR acts in part through Xrp1 as a downstream transcription factor. The DDR results in the repair of DNA damage or the removal of damaged cells by cell death. DNA repair can introduce errors (colored) including chromosome aberrations resulting in loss of genetic material (dashes) (C) When repair of DNA damage results in segmental aneuploidy, Xrp1 activation is triggered via an RpS12-dependent, p53-independent pathway that removes aneuploid cells via cell competition with unaffected cells.

### Xrp1 is an effector of p53 functions in the DDR

In imaginal discs, Xrp1 is induced by p53 activity in the acute phase of DDR at both RNA and protein levels (Figure 1). Unlike Xrp1 induction in *Rp^+/-^* cells, Xrp1 expression in the DDR occurs independently of autoregulatory transcription, which may be why the Xrp1-LacZ enhancer trap shows less upregulation than the endogenous Xrp1 transcript and protein. Multiple p53-dependent DDR genes, in fact, the majority in our study, were Xrp1-dependent, including the pro-apoptotic gene *hid*. Even though *hid-*reporters have been used as readouts of p53 activity (Fan et al., 2010; Zhang et al., 2014; Chakravarti et al., 2022; Ruiz-Losada et al., 2022), they may be more accurately described as targets of Xrp1. Other Xrp1 targets in the DDR include DNA repair genes, GstD1, and JNK signaling. Accordingly, Xrp1 is functionally required in the DDR. DNA damage-induced cell death is reduced in the absence of Xrp1, although not eliminated, perhaps because the pro-apoptotic gene *rpr* seems to be a direct target of p53, whose transcriptional induction does not require Xrp1. The absence of Xrp1 also leads to an increase in persistent DNA damage, as evidenced by persistent γH2Av staining. This is even more pronounced when p53 is absent as well (Figure 6). Persistent γH2Av staining may reflect ineffective DNA repair, insufficient apoptosis that permits badly damaged cells to survive, or both.

We show that in *Drosophila*, Xrp1 acts downstream to p53 as an intermediate transcription factor and probably controls the majority of DDR targets. So far, integrating data from gene expression studies, binding motif and ChIP-Seq searches, and global run-on sequencing of nascent mRNA has not yielded a consensus on the core set of direct p53 target genes in mammals (Riley et al., 2008; Allen et al., 2014; Fischer, 2017). It is possible that, as in *Drosophila*, mammalian p53 acts as a master transcription factor that regulates a network of gene expression responses both directly and through intermediate transcription factors.

### Xrp1 acts independently of p53 in cell competition

Although Xrp1 protein expression was almost completely p53-dependent during the acute DDR response, later Xrp1 expression was mostly p53-independent (Figure 6- figure supplement 1). Most DNA damage is normally resolved by the time of p53-independent Xrp1 expression (Wells et al., 2006). Without p53, however, DNA damage persisted longer than usual, so γH2Av labeling was still evident 24h after irradiation (Figure 6).

Previous work shows that p53-independent cell death can reflect cell competition of cells whose genomes have been altered by irradiation. Because 80 Rp are encoded by genes spread throughout the genome, a loss of 1-2% of the genome will often create haploinsufficiency for one or more Rp genes and trigger elimination by cell competition (Ji et al., 2021b). Consistent with this mechanism, the delayed, p53-independent Xrp1 expression largely depended on RpS12, a signal of defective ribosome biogenesis that is required to induce Xrp1 expression in *Rp^+/-^* cells and cells with defective rRNA synthesis (Lee et al., 2018b; Kiparaki et al., 2022). Accordingly, the p53-independent apoptosis occurring 24 hours after IR, and which is known to depend on RpS12, also depended on Xrp1. Xrp1 thus contributes to genome integrity at two levels, as a component of the p53-dependent DDR and as an executor of RpS12-dependent cell competition that functions, in part, to remove cells where DDR has failed or has altered cell genotypes (Figure 7B,C). Notably, Xrp1 expression levels were several-fold higher during the cell competition phase if p53 had been inhibited (Figure 6- figure supplement 1). Because this expression is p53-independent and likely to represent cell competition of aneuploid cells, it suggests that more aneuploid cells arise without p53 to coordinate the DDR. The persistence of H2Av labeling also suggested that double-strand breaks persist without p53. We think that without p53, DNA repair pathways, including DSB repair, single-strand break repair, and Nucleotide excision repair, are delayed or less effective, and coupled with faulty cell cycle arrest, result in a higher frequency of genome rearrangements and aneuploidy (Williams & Schumacher, 2016).

### Dual processes maintaining genome integrity following DNA damage

We have focused on two Xrp1-dependent processes contributing to genome integrity after DNA damage. The DDR depends on p53 in both *Drosophila* and mammals. Cell competition removes aneuploid cells in a p53-independent manner in *Drosophila* (Figure 7).

In mammals, the DDR was initially thought to represent the tumor suppressor function of p53. Later studies showed that tumor suppression is separable from the DDR and occurs afterwards (Christophorou et al., 2005; Christophorou et al., 2006; Li et al., 2012; Kaiser & Attardi, 2018). In mammals, where no clear Xrp1 homolog exists, Rp gene haploinsufficiency instead induces p53 through the nucleolar stress pathway (Bursac et al., 2014). Interestingly, differences in p53 activity levels between cells underlie multiple examples of cell competition in mammals (Bondar & Medzhitov, 2010b; Marusyk et al., 2010; Wagstaff et al., 2016; Fernandez-Antoran et al., 2019). Accordingly, we have suggested that in mammals, p53 takes on cell competition aspects of the Xrp1 function in *Drosophila* (Baker et al., 2019). Consistent with this, p53 mediates the removal of aneuploid cells in mammals, which Xrp1 does in *Drosophila* (Singla et al., 2020; Ji et al., 2021a). If p53 eliminates aneuploid cells in mammals by Rp gene dose-dependent cell competition, as Xrp1 does in *Drosophila*, then dual roles for mammalian p53 could both contribute to tumor suppression, as dual roles for Xrp1 maintain genome integrity in *Drosophila*. Consistent with this, analyses of early cancer development in a mouse pancreatic cancer model indicate that losses of single chromosomes or parts of chromosomes are very early steps in the progression toward transformation (Baslan et al., 2022). Mammalian cells with such small-scale monosomies only seem to survive in the absence of p53 (Chunduri et al., 2021). Although further studies will be required to confirm the suggestion, the observations are all consistent with the possibility that by contributing to genome integrity at the tissue level after the initial repair of DNA damage, cell competition could represent a significant tumor-suppressive process in mammals (Baker & Montagna, 2022).

## ACKNOWLEDGEMENTS

We thank Drs. J. Secombe, M. Kiparaki, S. Nair, A. Kumar, and S. Muliyil for comments on an earlier version of the manuscript. Fly stocks were obtained from the Bloomington Drosophila Stock Center (supported by NIH P40OD018537) and the Vienna Drosophila Resource Center. Confocal microscopy was performed in the Analytical Imaging Facility of the Albert Einstein College of Medicine (supported by the NCI P30CA013330) using the Leica SP8 microscope acquired through NIH SIG 1S10 OD023591. The monoclonal antibody mAb40-1a developed by J. R. Sanes was obtained from the Developmental Studies Hybridoma Bank, created by the NICHD and maintained at the University of Iowa, Department of Biology, Iowa City, IA 52242. This project was supported by a grant from the National Institutes of Health (R01-CA284362 to NEB). Part of this work is supported by the Division of Intramural Research at the National Heart, Lung, and Blood Institute within the National Institutes of Health (1ZIAHL006126 to NMR).

## Disclaimer

Part of this research was supported by the Intramural Research Program of the National Institutes of Health (NIH). The contributions of the NIH authors were made as part of their official duties as NIH federal employees, are in compliance with agency policy requirements, and are considered Works of the United States Government. However, the findings and conclusions presented in this paper are those of the authors and do not necessarily reflect the views of the NIH or the U.S. Department of Health and Human Services.

## MATERIAL AND METHODS

### 1. Fly husbandry

Flies were raised at 25°C on standard cornmeal and molasses medium, containing the following ingredients per 1L: 18g yeast; 22g molasses; 80g malt extract; 9g agar; 65g cornmeal; 2.3g methyl para-benzoic acid; 6.35ml propionic acid. Larvae were not distinguished for their sex before dissection.

### 2. DNA damage γ-radiation treatment

Flies were grown in the standard cornmeal medium at 25°C. After egg laying was done for 24 hours, the flies were transferred to a fresh vial. Following this, the larvae were allowed to grow until the wandering late III instar stage, and DNA damage was induced by treating them with γ-radiation from cesium-137 using a Shepherd irradiator. For most experiments, 1500 Rads were administered, and dissection was performed at a specified time point after the irradiation.

### 3. Cell death quantification

Cell death was quantified by counting the number of Dcp-1 positive cells with the help of the ImageJ cell counter plugin. For UAS-Gal4 experiments, Dcp-1 positive cells in Gal4 to the control compartment were counted and compared as the ratio across different genetic perturbations. In the case of mutant wing discs, the number of Dcp-1 positive cells was counted in the entire disc, and the absolute number was compared.

### 4. Immunohistochemistry

Imaginal discs were fixed using 4% formaldehyde in 1× PBS, followed by 2 h blocking with 0.5% bovine serum albumin (BSA) dissolved in 0.1% PBST (1×PBS with 0.1% Triton X-100). Primary antibody incubations were performed at 4°C overnight, followed by 2 hours of secondary incubation at room temperature. All washes following antibody incubation were performed in 0.1% PBST. Primary antibodies were against rabbit anti-cleaved Dcp-1 (Cell Signaling 9578, 1:200), mouse anti-γH2Av (DSHB UNC93-5.2.1, 1:50), mouse anti-βGalactosidase (DSHB 40a-1, 1:50), mouse anti-p53 (DSHB 25F4, 1:50), and mouse anti-GFP (DSHB 12A6, 1:50), mouse anti-HA (CST 2367, 1:100), and anti-Xrp1 (1:250; gift from Rio lab). Alexa-Fluor (Thermo Fisher-Invitrogen) and Cyanine dye (Jackson labs) conjugated secondary antibodies were used at 1:200 dilution. Image acquisition was performed on the Leica-Sp8 microscope, intensity adjustments were made using Photoshop CC 2018, and Image J was used for image processing.

Previous experiments have supported that significant results could be obtained from 5 replicates; in most cases, more than 5 samples were analyzed. Experiments were repeated at least 2 times or more for most of the results. No exclusion of the data points was made while performing statistical significance. For the quantitative data, N values are indicated in the figure legends. Statistical significance was performed using a Student’s t-test for independent or dependent means, as shown in the figure legends. For specific data sets, one-way ANOVA with Post Hoc Turkey HSD was performed, as mentioned in the figure legends. Exact p-values are indicated in the figures.

### 5. q-RT PCR

Total RNA was extracted from wing imaginal discs of late-third instar larvae treated with 2000 Rads of IR; untreated larvae were taken as a control for normalization. Wing discs were collected in TRIzol reagents (Ambion), and RNA was isolated according to the manufacturer’s instructions. RNA samples were DNase-treated according to the manufacturer’s instructions (Thermo Fisher; AM1907) and reverse transcribed using the Verso cDNA Synthesis Kit (Thermo Fisher; AB1453). The cDNA was used to perform the q-RT PCR reactions following the manufacturer’s instructions for the TaqMan^TM^ Fast Advanced Master Mix (Thermo Fisher; 4444557). Custom design FAM-MGB TaqMan probes were utilized for q-RT-PCR. The gene expression values were normalized to the housekeeping gene RpL32 and confirmed with actin as a second control (data not included). The following calculations were used for estimating fold change in gene expression Δ: Ct = Ct (gene of interest) – Ct (housekeeping gene), ΔΔCt = ΔCt (3-4 hr IR) – ΔCt (untreated average), and Fold gene expression = 2^ -(ΔΔCt). 3 biological replicates were analysed for each condition and genotype. The gene expression analysis was performed using technical triplicates for each sample. It has been previously established that significant results can obtained 3 biological and 3 technical triplicates.

The following TaqMan probes were used in this study:

1. p53 Dm02154335_g1
2. Xrp1 Dm02142088_g1
3. rpr Dm01823054_s1
4. hid Dm01823031_m1
5. GstD1 Dm02362792_s1
6. eiger Dm01794373_m1
7. rad50 Dm01817359_g1
8. PolZ1 Dm01843262_g1
9. DNAlig4 Dm01838632_g1
10. Ku80 Dm01791676_g1
11. *Drosophila* RpL32 Dm02151827_g1
12. *Drosophila* actin Dm02361909_s1

---

**Figure 1- Supplement 1:**
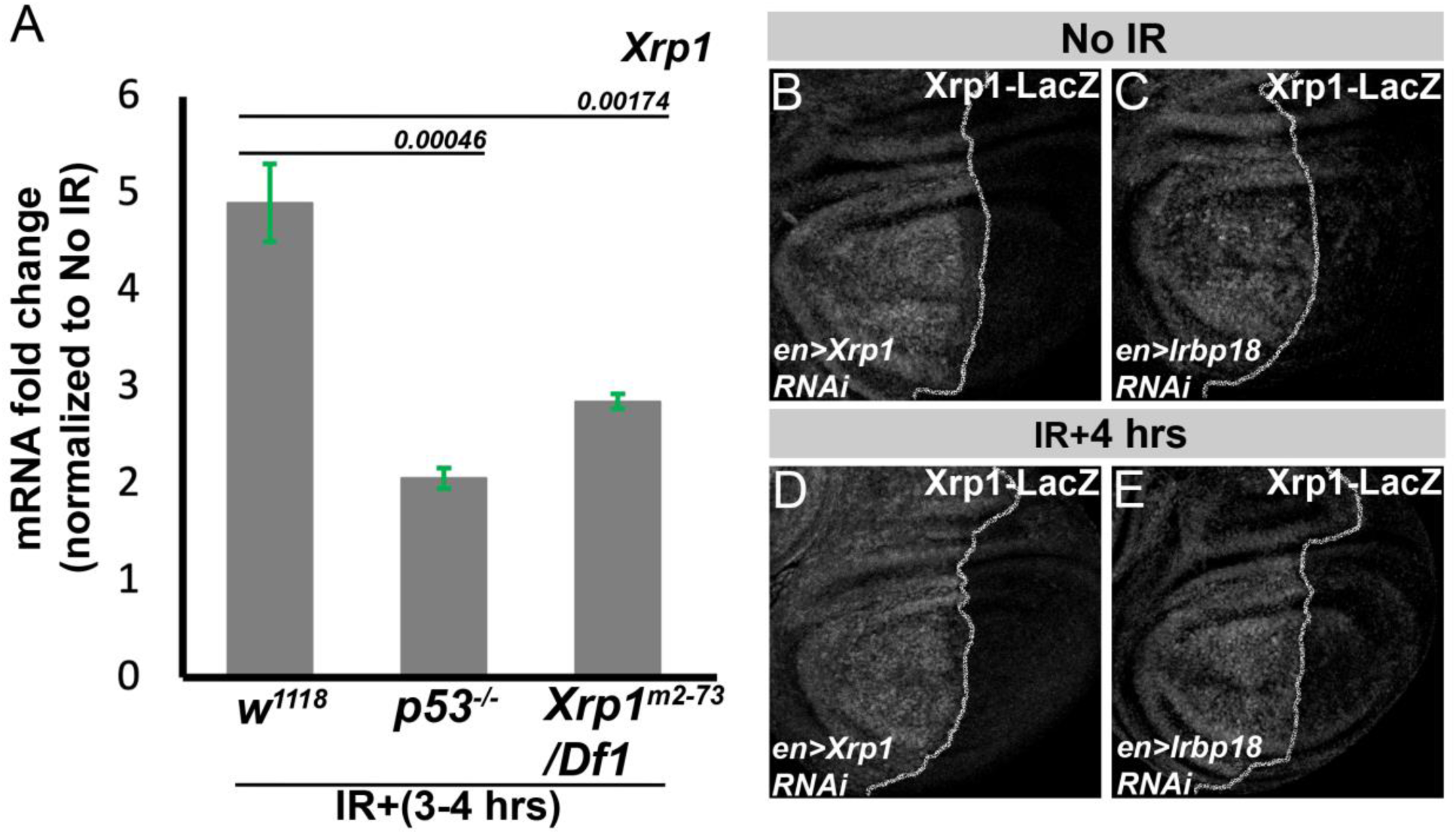
Xrp1 RNA expression in response to DNA damage. qRT-PCR analysis for Xrp1 mRNA expression levels from III instar wing discs from the genotypes *w^1118^*, *p53^[5A-1-4]^*, and *Xrp1^m2-73^/Df1* collected 3-4 hours after irradiation (1500 RADs) and normalized to an untreated (0 RADs) sample (A). III instar wing discs assayed for *Xrp1^02515^*-lacZ enhancer trap expression by anti-βgal antibody in 0 Rads for *en>Xrp1 RNAi* (B), *en>Irbp18 RNAi* (C), and 4 hours after 1500 Rads treatment in genotypes *en>Xrp1 RNAi* (D), *en>Irbp18 RNAi* (E). Number of wing discs analyzed for genotypes was N=3 for B, N=2 for C, N=3 for D, and N=5 for E Data is plotted as mean± SEM, and a two-tailed student t-test for independent means was performed for statistical significance. Genotype: (A) *w^1118^*, *yw; p53^[5A-1-4]^,* and *Xrp1^m2-73^/Df 1*, (B & C) *en-Gal4::UAS-RFP, UAS-Xrp1 RNAi* (VDRC 107860); *P{PZ}Xrp1^02515^* (C & E) *en-Gal4::UAS-RFP, UAS-Irbp18 RNAi*; *P{PZ}Xrp1^02515^*

**Figure 3- Supplement:**
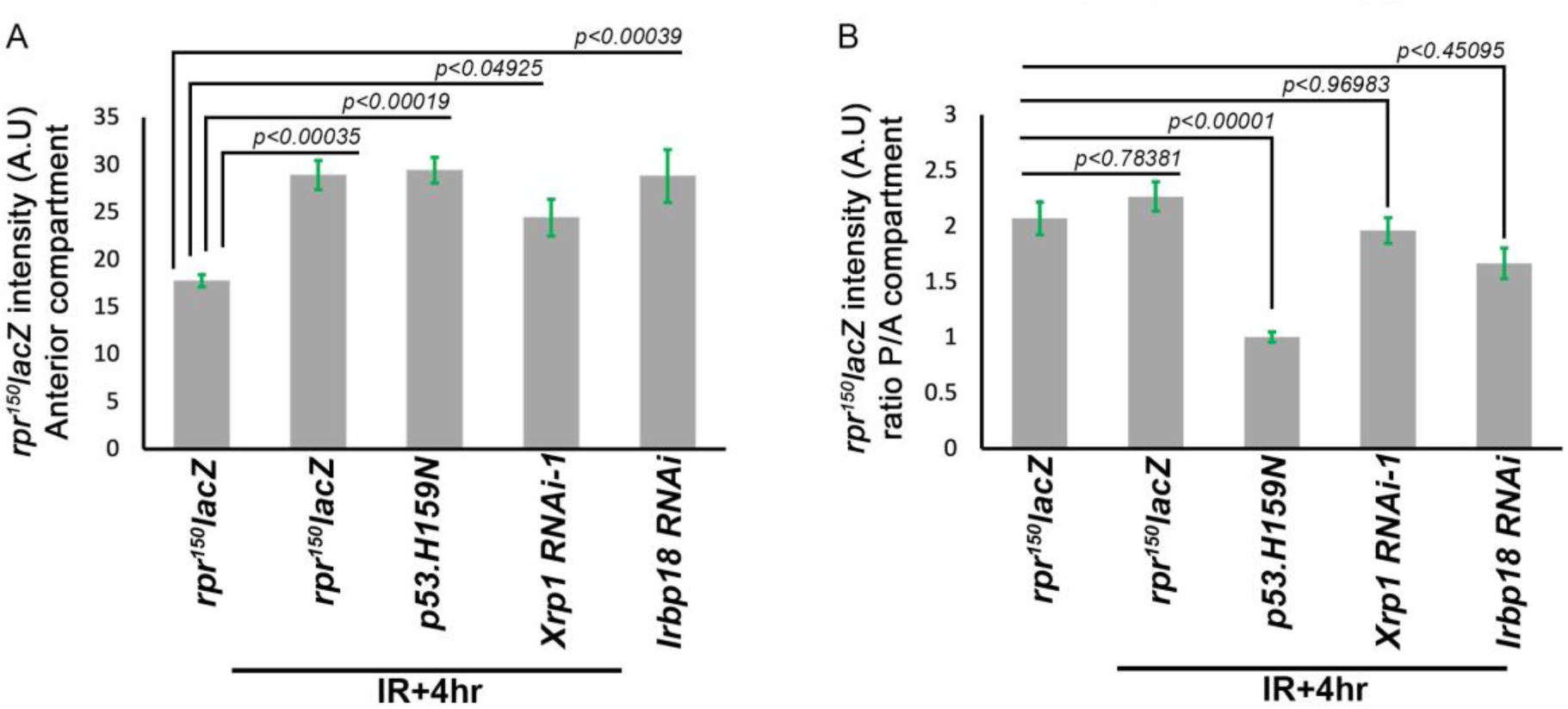
*rpr^150^lacZ* intensity quantification. (A) Average intensity quantification for *rpr^150^lacZ* from the wing, the anterior compartment of the wing pouch, with and without IR for the genotypes shown in Figure 3 L-P. *rpr^150^lacZ* average intensity in the anterior compartments is elevated by IR, regardless of gene expression or depletion in the P compartments. (B) The mean posterior (P) to mean anterior (A) LacZ expression ratio for each was only reduced by p53-DN, which decreased the relative *rpr*-*LacZ* expression level. Statistics: Data is plotted as mean± SEM; n=5 for *irbp18* RNAi, n=7 for other genotypes; significance assessed by one-way ANOVA with Post Hoc Turkey HSD. For panel B, genotypes were compared only to the unirradiated control. Genotype: (A & B) *rpr^150^*-lacZ, *rpr^150^-lacZ, UAS-p53^H159N^* (Bl. 8420)*; hh-Gal4::UAS-GFP*, *rpr^150^-lacZ, UAS-Xrp1 RNAi* (VDRC 107860)*; hh-Gal4::UAS-GFP, rpr^150^-lacZ, UAS-Irbp18 RNAi; hh-Gal4::UAS-GFP*

**Figure 4- Supplement 1:**
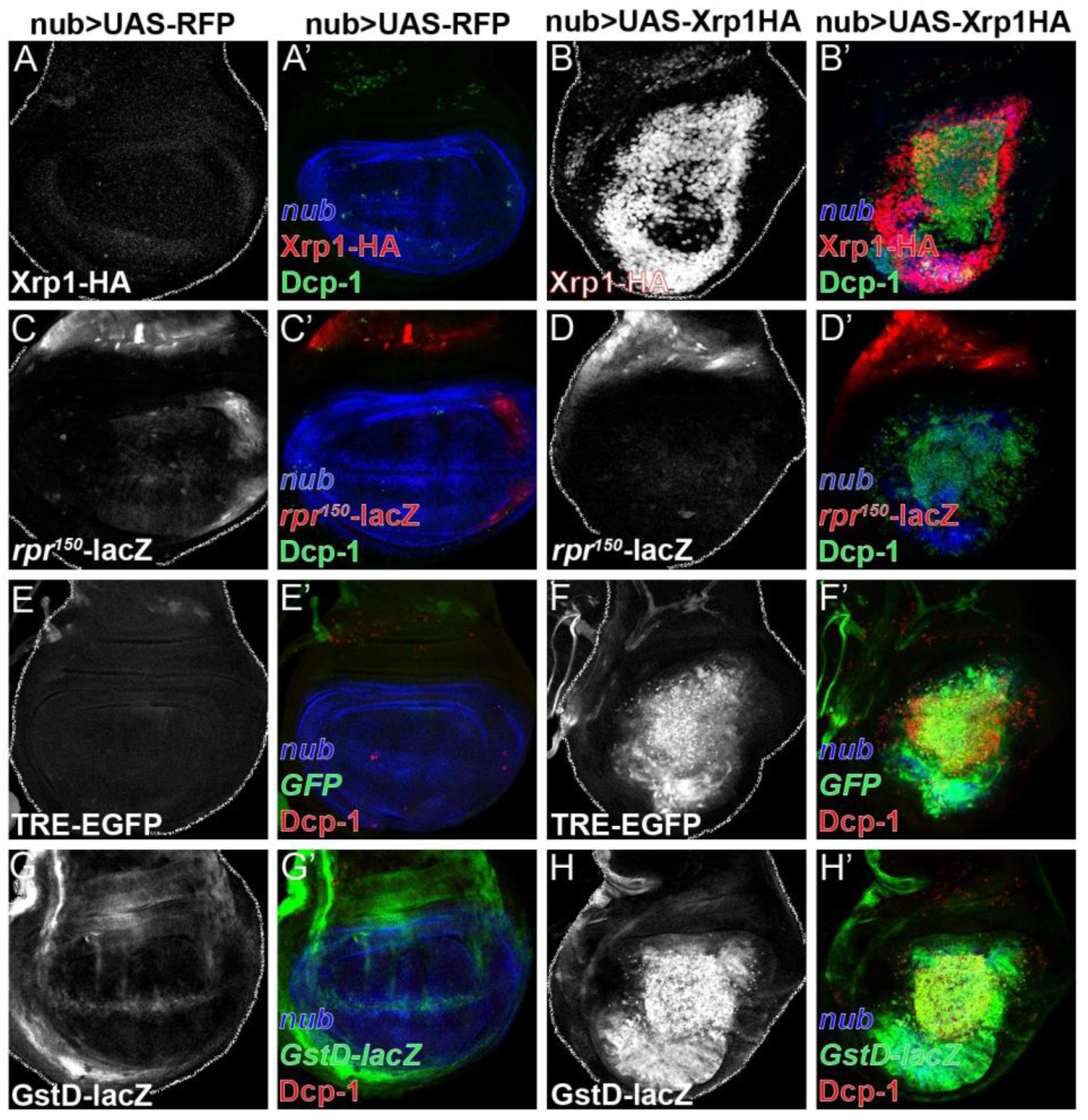
Xrp1 is sufficient for the expression of p53-dependent DDR target genes. Xrp1-HA protein measured by anti-HA (red) and cell death by Dcp-1 (green) in the III instar wing disc expressing UAS-RFP control (A) and UAS-Xrp1 long isoform (B) under the control of *nub-Gal4* (blue). *rpr^150^-lacZ* reporter expression measured by anti-βgal (green) and cell death by anti-Dcp-1 (red) in III instar wing imaginal disc expressing UAS-RFP (C) and UAS-Xrp1 (D) under the control of *nub-Gal4*. *TRE-EGFP* reporter expression measured by anti-GFP (green) and cell death by anti-Dcp-1 (red) in III instar wing imaginal disc expressing UAS-RFP control (E) and UAS-Xrp1 (F) under the control of *nub-Gal4* (blue). *GstD-lacZ* reporter expression measured by anti-GFP (green) and cell death by anti-Dcp-1 (red) in III instar wing imaginal disc expressing UAS-RFP (G) and UAS-Xrp1 (H) under the control of *nub-Gal4* (blue). N=7-8 discs were analyzed for every genotype, and each experiment was repeated at least 2 times. All the control and experimental genotypes shown in this figure were kept at 18°C to lessen the effect of Xrp1 overexpression. Genotype: (A) *nub-Gal4::UAS-RFP,* (B) *nub-Gal4::UAS-RFP::UAS-Xrp1-HA,* (C) *nub-Gal4::UAS-RFP, rpr^150^-lacZ,* (D) *nub-Gal4::UAS-RFP::UAS-Xrp1-HA, rpr^150^-lacZ,* (E) *nub-Gal4::UAS-RFP, TRE-GFP,* (F) *nub-Gal4::UAS-RFP::UAS-Xrp1-HA, TRE-GFP,* (G) *nub-Gal4::UAS-RFP, GstD-lacZ,* (H) *nub-Gal4::UAS-RFP::UAS-Xrp1-HA, GstD-lacZ*

**Figure 5- Supplement:**
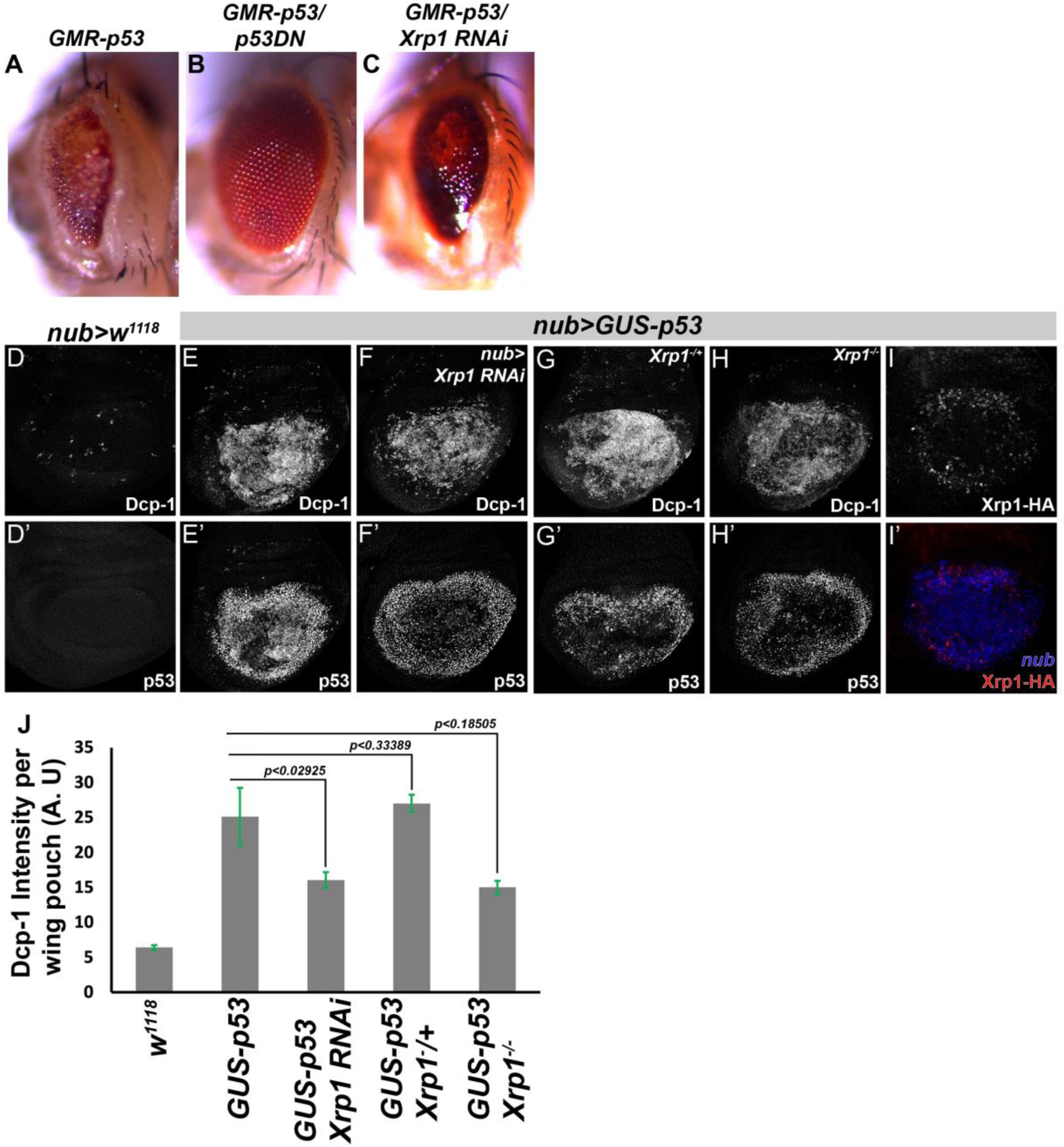
Xrp1 do not rescue p53 O/E phenotype. (A) Bright field photograph of *Drosophila* eye, rough eye phenotype due to constitutive expression of p53 under the control of GMR-promoter (*GMR-p53*). (B) Overexpression of dominant-negative p53 (*UAS-p53^H159N^*) rescues the rough eye phenotype. (C) Xrp1 knockdown by RNAi has a slight effect on the rough eye phenotype. (D) Control *nub-Gal4* exhibits few Dcp-1 positive cells, and no p53 protein was detected by anti-p53 antibody. (E) *nub-Gal4* driving p53 overexpression (*GUS-p53*) in the wing pouch exhibits elevated cell death measured by Dcp-1 and p53 protein, measured by anti-p53 antibody. (F) Co-expressing *UAS-Xrp1* RNAi and *GUS-p53* under *nub-Gal4* led to a decrease in cell death by approximately 50%. (G) Removing one copy by *Xrp1^m2-73^/+* ^did not affect^ cell death. (H) Removing both copies of Xrp1 (*Xrp1^m2-73^/Xrp1^attp flox^*) results in a decrease in cell death of approximately 50%. Xrp1-HA expression was assayed by an anti-HA antibody in the wing disc, expressing p53 under the control of *nub-Gal4* (I & I’). (J) Dcp1 intensity was quantified using ImageJ; N = 6 for each genotype. Data is plotted as mean± SEM, two-tailed student t-test for independent means. All the control and experimental genotypes shown in this figure were kept at 18°C to lessen the effect of p53 overexpression. Genotype: (A) *GMR-p53,* (B) *GUS-p53, GMR-Gal4; UAS-p53^H159N^* (Bl. 4021), (C) *GUS-p53, GMR-Gal4; UAS-Xrp1 RNAi* (Bl. 4521), (D) *nub-Gal4; w^1118^*, (E) *nub-Gal4, GUS-p53,* (F) *nub-Gal4, GUS-p53; UAS-Xrp1 RNAi* (Bl. 4521), (G) *nub-Gal4, GUS-p53; Xrp1^m2-73^/+,* (H) *nub-Gal4, GUS-p53; Xrp1^m2-73^/Xrp1^attp flox^,* (I) *nub-Gal4, GUS-p53; Xrp1-HA*

**Figure 6- Supplement 1:**
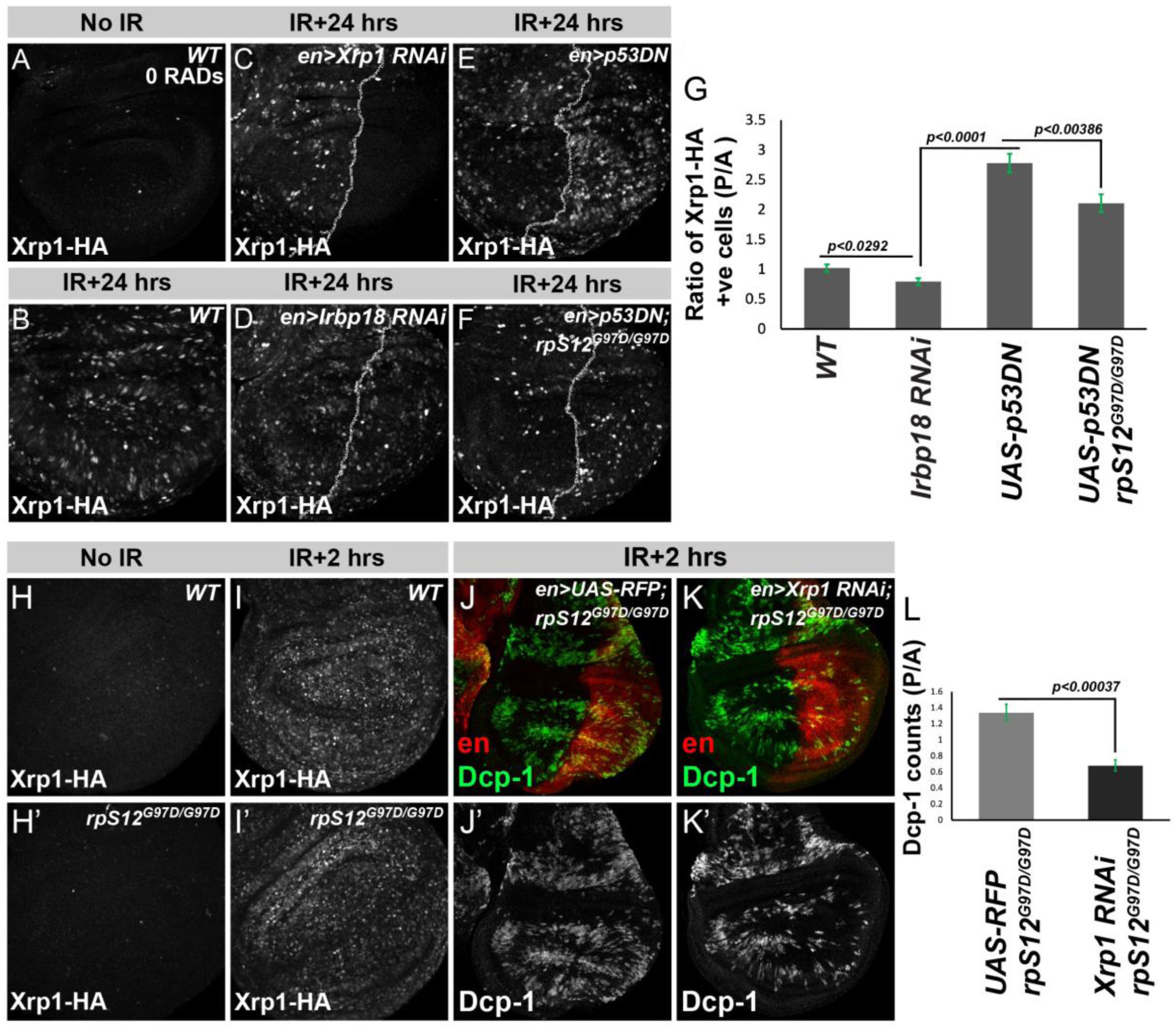
Xrp1 induction independent of p53. (A-F) Xrp1-HA expression in control disc (A), 24 hours after 1500 rads (B), *UAS-Xrp1* RNAi (C), *UAS-Irbp1* RNAi (D), *UAS-p53^H159N^* (E), and *UAS-p53^H159N^* in *rpS12^G97D/G97D^* mutant background (F). (G) The ratio of Xrp1-positive cells in anterior (A) to posterior (P) compartments in *UAS-p53^H159N^* and *UAS-p53^H159N^* in *rpS12^G97D/G97D^* mutant background. (H-J) Cell death measured by anti-Dcp-1 (green and grayscale) in wing imaginal discs dissected at 2 hours after 1500 Rads, measured in en-Gal4 (posterior compartment (red)) versus anterior compartment *UAS-RFP; rpS12^G97D^* (H), and *UAS-Xrp1 RNAi; rpS12^G97D^* (I). Dcp-1 positive cells were quantified in anterior (A) and posterior (P) compartments and plotted as the P/A ratio (J). N=7 discs for (A, B, C, E, and F), N=3 discs for (D), N=6 discs for (J-L), data is plotted as mean± SEM, and a student t-test, two-tailed for the independent sample, was performed for statistical significance. Genotype: (A & B) *Xrp1-HA*, (C) *en-Gal4::UAS-RFP; UAS-Xrp1 RNAi* (VDRC 107860)*; Xrp1-HA*, (D) *en-Gal4::UAS-RFP; UAS-Irbp18 RNAi; Xrp1-HA*, (E) *en-Gal4::UAS-RFP; UAS-p53DN^H159N^* (Bl. 8420)*; Xrp1-HA*, (F) *en-Gal4::UAS-RFP; UAS-p53DN^H159N^* (Bl. 8420)*; Xrp1-HA-rpS12^G97D/G97D^* (H) *en-Gal4::UAS-RFP; rpS12^G97D/G97D^*, (I) *en-Gal4::UAS-RFP, UAS-Xrp1 RNAi* (VDRC 107860)*; rpS12^G97D/G97D^*

**Figure 6- Supplement 2:**
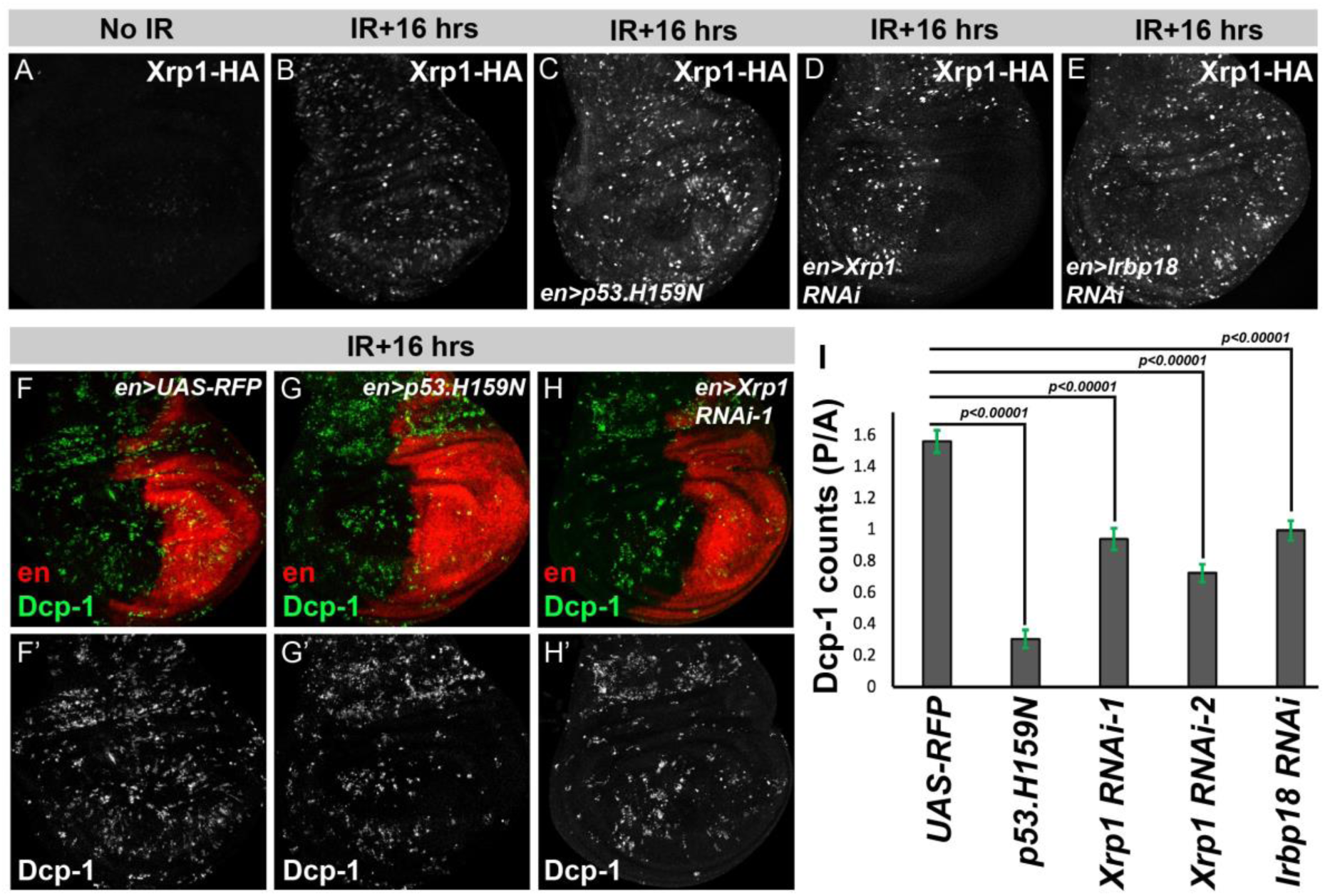
Induction of p53-independent DDR occurs around 16 hours post-irradiation. (A-E) Xrp1-HA expression in control disc (A), 16 hours after 1500 rads (B), *UAS-p53^H159N^* (C), *UAS-Xrp1* RNAi (D), and *UAS-Irbp1* RNAi (E). *Drosophila* III instar wing imaginal discs dissected at 16 hours after 1500 Rads, cell death is measured by Dcp-1 (green and grayscale) in different genetic perturbations using en-Gal4 (posterior compartment (red)), UAS-RFP (F), *UAS-p53^H159N^* (G), and *UAS-Xrp1* RNAi (H). Dcp-1 positive cells were quantified and plotted as the ratio of Dcp-1 positive cells in posterior (P) to anterior (A) compartments (I). Dcp-1 positive cells were quantified in anterior (A) and posterior (P) compartments and plotted as the P/A ratio (I). N=10 for (F-I), data is plotted as mean± SEM, and a student t-test, two-tailed for the independent sample, was performed for statistical significance. Genotype: (A & B) *Xrp1-HA*, (C) *en-Gal4::UAS-RFP; UAS-p53DN^H159N^* (Bl. 8420)*; Xrp1-HA*, (D) *en-Gal4::UAS-RFP; UAS-Irbp18 RNAi; Xrp1-HA*, and (E) *en-Gal4::UAS-RFP; UAS-Xrp1 RNAi* (VDRC 107860)*; Xrp1-HA*, (F) *en-Gal4::UAS-RFP,* (G) *en-Gal4::UAS-RFP, UAS-p53DN^H159N^* (Bl. 8420), (H) *en-Gal4::UAS-RFP, UAS-Xrp1 RNAi* (Bl. 4521), (I) *en-Gal4::UAS-RFP, UAS-Irbp18 RNAi*

**Supplementary Table 1:** Key Resources Table.

| **Key Resources Table** |  |  |  |  |
| --- | --- | --- | --- | --- |
| **Reagent type (species) or resource** | **Designation** | **Source or reference** | **Identifiers** | **Additional information** |
| gene (*Drosophila melanogaster*) | Xrp1 |  | FLYBASE: FBgn0261113 |  |
| gene (*Drosophila melanogaster*) | irbp18 |  | FLYBASE: FBgn0036126 |  |
| genetic reagent (*D. melanogaster*) | p53 |  | FLYBASE: FBgn0039044 |  |
| genetic reagent (*D. melanogaster*) | rpS12 |  | FLYBASE: FBgn0286213 |  |
| genetic reagent (*D. melanogaster*) | *Xrp1*<sup>m2-73</sup> | (Lee et al., 2016) | FLYBASE: FBal0346068 | Bloomington Drosophila Stock Center #81270 |
| genetic reagent (*D. melanogaster*) | *Xrp1*<sup>02515</sup> | (Spradling et al., 1999) | FLYBASE: FBal0009448 | Bloomington Drosophila Stock Center #11569 |
| genetic reagent (*D. melanogaster*) | *Xrp1*<sup>attP</sup><sub>flox</sub> | (Blanco et al., 2020) |  | Strain maintained in Dr. Nicholas Baker's lab |
| genetic reagent (*D. melanogaster*) | Xrp1<sup>HA</sup> | (Blanco et al., 2020) |  | Strain maintained in Dr. Nicholas Baker's lab |
| genetic reagent (*D. melanogaster*) | UAS-Xrp1 (long isoform) | (Tsurui-Nishimura et al., 2013) | FLYBASE: FBal0284387 |  |
| genetic reagent (*D. melanogaster*) | *irbp18*<sup>05006</sup> | (Francis et al., 2016) | FLYBASE: FBal0185260 | Exelisis Drosophila Collection #f05006 |
| genetic reagent (*D. melanogaster*) | *rpS12*<sup>G97D</sup> |  | FLYBASE: FBgn0286213 | Strain maintained in Dr. Nicholas Baker's lab |
| genetic reagent (*D. melanogaster*) | *p53*<sup>[5A-1-4]</sup> |  | FLYBASE: FBgn0039044 | Bloomington Drosophila Stock Center #6815 |
| genetic reagent (*D. melanogaster*) | UAS-p53.H15 9N.Ex-2 (Chr. II) |  | FLYBASE: FBti0040583 | Bloomington Drosophila Stock Center #8420 |
| genetic reagent (*D. melanogaster*) | UAS-p53.H15 9N.Ex -3 (Chr. III) |  | FLYBASE: FBti0040584 | Bloomington Drosophila Stock Center #8421 |
| genetic reagent (*D. melanogaster*) | UAS-dsRNA<sup>Xr p1</sup> | (Ni et al., 2011) | FLYBASE: FBti0144696 | Bloomington Drosophila Stock Center #34521 |
| genetic reagent (*D. melanogaster*) | UAS-dsRNA<sup>Xr</sup><sub>p1</sub> |  | FLYBASE: FBst0479673 | Vienna Drosophila Resource Center #107860 |
| genetic reagent (*D. melanogaster*) | UAS-dsRNA<sup>irb</sup><sub>p18</sub> | (Ni et al., 2011) | FLYBASE: FBti0140125 | Bloomington Drosophila Stock Center #33652 |
| genetic reagent (*D. melanogaster*) | UAS-dsRNA<sup>w</sup> | (Ni et al., 2011) | FLYBASE: FBtp0064645 | Bloomington Drosophila Stock Center #33623 |
| genetic reagent (*D. melanogaster*) | arm-LacZ | (Vincent et al., 1994) | FLYBASE: FBal0040819 |  |
| genetic reagent (*D. melanogaster*) | Ubi-GFP | (Davis et al., 1995) | FLYBASE: FBal0047085 |  |
| genetic reagent (*D. melanogaster*) | M(2)56F (mutating RpS18) | Laboratory of Y. Hiromi | FLYBASE: FBal0011916 |  |
| genetic reagent (*D. melanogaster*) | hs-FLP | (Struhl & Basler, 1993) | FLYBASE: FBtp0001101 |  |
| genetic reagent (*D. melanogaster*) | ey-FLP | (Newsome et al., 2000) | FLYBASE: FBal0098303 |  |
| genetic reagent (*D. melanogaster*) | GMR-gal4 | (Freeman, 1996) | FLYBASE: FBtp0001315 |  |
| genetic reagent (*D. melanogaster*) | P{neoFR T}42D | (Xu & Rubin, 1993) | FLYBASE: FBti0141188 | Bloomington Drosophila Stock Center #1802 |
| genetic reagent (*D. melanogaster*) | P{neoFR T}80B | (Xu & Rubin, 1993) | FLYBASE: FBti0002073 | Bloomington Drosophila Stock Center #1988 |
| genetic reagent (*D. melanogaster*) | P{neoFR T}82B | (Xu & Rubin, 1993) | FLYBASE: FBti0002074 | Bloomington Drosophila Stock Center #2050, 2051 |
| Antibody | Polyclonal Rabbit anti-XRP1 (short isoform) | (Francis et al., 2016) |  | 1:200 dilution |
| Antibody | Mouse anti-HA Tag monoclonal | Cell Signalling Technology | Cat #2367<br>RRID:AB_10691311 | 1:100 dilution |
| Antibody | Mouse anti-beta galactosidase (mAb40-1a) monoclonal | DSHB | RRID: AB_2314509 | 1:100 dilution |
| Antibody | Rabbit anti-active-Dcp1 polyclonal | Cell Signalling Technology | Cat #9578<br>RRID:AB_2721060 | 1:50 dilution |
| Antibody | Mouse anti-p53 (25F4) monoclonal | DSHB | RRID: AB_579787 | 1:20 dilution |

